# Therapeutic vulnerability to PARP1/2 inhibition in *RB1*-mutant osteosarcoma

**DOI:** 10.1101/2020.12.28.424497

**Authors:** Georgia Zoumpoulidou, Carlos A Mendoza, Caterina Mancusi, Ritika M Ahmed, Milly Denman, Christopher D Steele, Jiten Manji, Nischalan Pillay, Sandra J Strauss, Sibylle Mittnacht

## Abstract

**Background:** Loss-of-function mutations of the retinoblastoma tumour suppressor *RB1* are key drivers in cancer, with prominent involvement in the natural history of Osteosarcoma (OS). *RB1* loss-of-function compromises genome maintenance in cells and hence could yield vulnerability to therapeutics targeting such processes.

**Method:** We assessed the response to Poly-ADP-Polymerase1/2 inhibitors (PARPi) in histiotype-matched cancer cell lines differing in *RB1* status including an extended panel of OS lines, measuring viability, clonogenic activity and inhibition of xenograft growth *in vivo*. We used mutational signature analysis and RAD51 immunostaining to assess competence for homologous repair defect (HRd).

**Results:** We report selective hypersensitivity to clinically-approved PARPi in OS lines with RB1 mutation, which extends to other cancer histiotypes and is induced in RB1-normal OS following engineered RB1 loss. PARPi treatment caused extensive cell death in RB1-mutated OS and extended survival of mice carrying human RB1-mutated OS grafts. Sensitivity in OS with natural or engineered RB1 loss surpassed that seen in BRCA-mutated backgrounds where PARPi are showing clinical benefit. PARPi sensitivity was not associated with loss of RAD51 recruitment and HRd-linked mutational signatures, which predict PARPi sensitivity in cancers with BRCA1/2 loss, but linked to rapid activation of replication checkpoint signalling with S phase transit critical for the death response observed.

**Conclusion:** Our work demonstrates that mutations in *RB1* causes clinically relevant hypersensitivity to approved PARP1/2-targeting therapeutics and advocates PARP1/2 inhibition as a novel, genome lead strategy for *RB1*-mutated osteosarcoma.

## INTRODUCTION

Biallelic mutations targeting the tumour suppressor RB1 are prominently associated with difficult to treat cancers, including osteosarcoma.

Osteosarcoma or osteogenic sarcoma, is the most common primary human malignancy in bone. More than half of cases arise in children and young adults and disproportionately contribute to cancer death in these age groups [1]. Aggressive multimodal treatment involving combination chemotherapy has substantially increased survival in this disease. However, less than 30% of patients diagnosed with metastatic disease show long term response, and relapse or treatment associated toxicity in patients diagnosed with localised disease remain chief concerns [2], [3], [1], [4].

Emerging osteosarcoma genomics data is revealing the prominent presence of deleterious mutations in the known tumour suppressors *TP53, RB1, RECQL4, BLM, and WRN. RB1* mutations are seen in 40-60% of sporadic osteosarcoma [5], [6], [7] making it the second most frequently mutated gene in this disease after *TP53*. Studies of osteosarcoma genomic evolution invariably report RB1 mutations as early, truncal events [8], [9] and germline mutations in *RB1* increase the risk of osteosarcoma development [10] supporting a causal role of *RB1* defects in disease initiation. Notably, various sources, including a recent systematic review, report association of *RB1* mutation with poor prognosis including an increased risk of metastasis [11], paralleling observations in other cancers with *RB1* involvement [12], [13] and indicating a clear unmet clinical need in patients with *RB1*-mutant osteosarcoma.

Currently the majority of osteosarcoma patients are treated in an identical manner, irrespective of presentation or genotype [3]. While conventional combination chemotherapy has remained standard of care for osteosarcoma, targeted agents including multitargeted tyrosine kinase inhibitors are showing efficacy in early phase clinical trials albeit with significant grade 3 to 5 toxicity liabilities [14], and may provide additional options in relapsed disease. Personalised, biomarker-informed treatments have been proposed in preclinical work for various gain of function events [5], [7], [15], [5], indicating targeted, genome-informed treatment could provide future solutions in osteosarcoma. However, opportunities identified by the highly prevalent loss-of-function events, including *TP53* and *RB1*, have not been reported.

*RB1* is a negative regulator of the cell cycle but has been ascribed other functions [16]. RB1 defects in cells cause complex changes in cell response including anomalies in DNA double-strand break (DDSB) repair [17], [18], [19] and mitotic fidelity [20]. Such DNA metabolic alterations raise the possibility that synthetic lethal opportunities may exist involving therapeutics known to interact with defective repair and mitosis.

Based on assessments involving an extensive osteosarcoma focused cell line panel, we here report selective sensitivity of *RB1*-mutant osteosarcoma to inhibitors of Poly-(ADP-Ribose)-Polymerase1,2 (PARPi). PARP1,2 enzymes have complex roles in single-strand-break DNA repair, transcription and replication [21]. PARP inhibition is selectively lethal in cancers cells with mutation in the BRCA1/2 tumour suppressors causing defective homologous recombination (HRd) [22], and multiple PARPi have regulator-approval for 2^nd^ and 1^st^ line treatment in HRd and/or BRCA1/2 mutated ovarian, breast and pancreatic cancers and recent FDA breakthrough status in castration resistant prostate cancer [23], [24].

We document highly penetrant PARPi hypersensitivity following from *RB1* mutation, with dose-sensitivity similar to that caused by BRCA1/2 mutation. We validate the involvement of *RB1* defects in this response and document single-agent PARPi efficacy in a preclinical model of RB*1*-mutant osteosarcoma. Our work proposes a novel genome led strategy for treatment of osteosarcoma, involving stratified use of PARP1/2 targeting therapeutics.

## RESULTS

### Differential PARP1/2 inhibitor sensitivity in *RB1*-mutant osteosarcoma tumour cell lines

To identify therapeutically exploitable vulnerability linked to deleterious *RB1* mutation we assessed the sensitivity of histiotype-matched cancer cell pairs differing in *RB1* mutation status focussing on approved clinical agents that target DNA metabolic processes.

Day-5 viability assessments using resazurin reduction revealed consistent hypersensitivity to the PARPi olaparib in *RB1*-mutant compared to matched *RB1*-normal lines (Supplementary Figure 1A-C). A strong association between olaparib sensitivity and *RB1* status extended to a poly-cancer cell line panel, with median area-under-the-curve (AUC) values significantly lower in *RB1*-mutant compared to *RB1*-normal lines (Supplementary Figure 1D-F), indicative that *RB1* status in cancers is associated with, and may predict hypersensitivity to PARPi.

To examine whether this selective sensitivity extends to osteosarcoma we measured the response to olaparib across a broad osteosarcoma-focussed cell panel. To benchmark clinically relevant response we included the pancreatic cancer cell line Capan-1, known for profound PARPi sensitivity due to defective *BRCA2* [25]. This assessment confirmed increased dose sensitivity (Figure 1A), yielding a highly significant differential in median sensitivity, assessed using AUC values (Figure 1B), in osteosarcoma lines with known *RB1*-mutant status and/or lacking detectable RB1 expression (Figure 1L) compared to *RB1*-normal lines.

**Figure 1:**
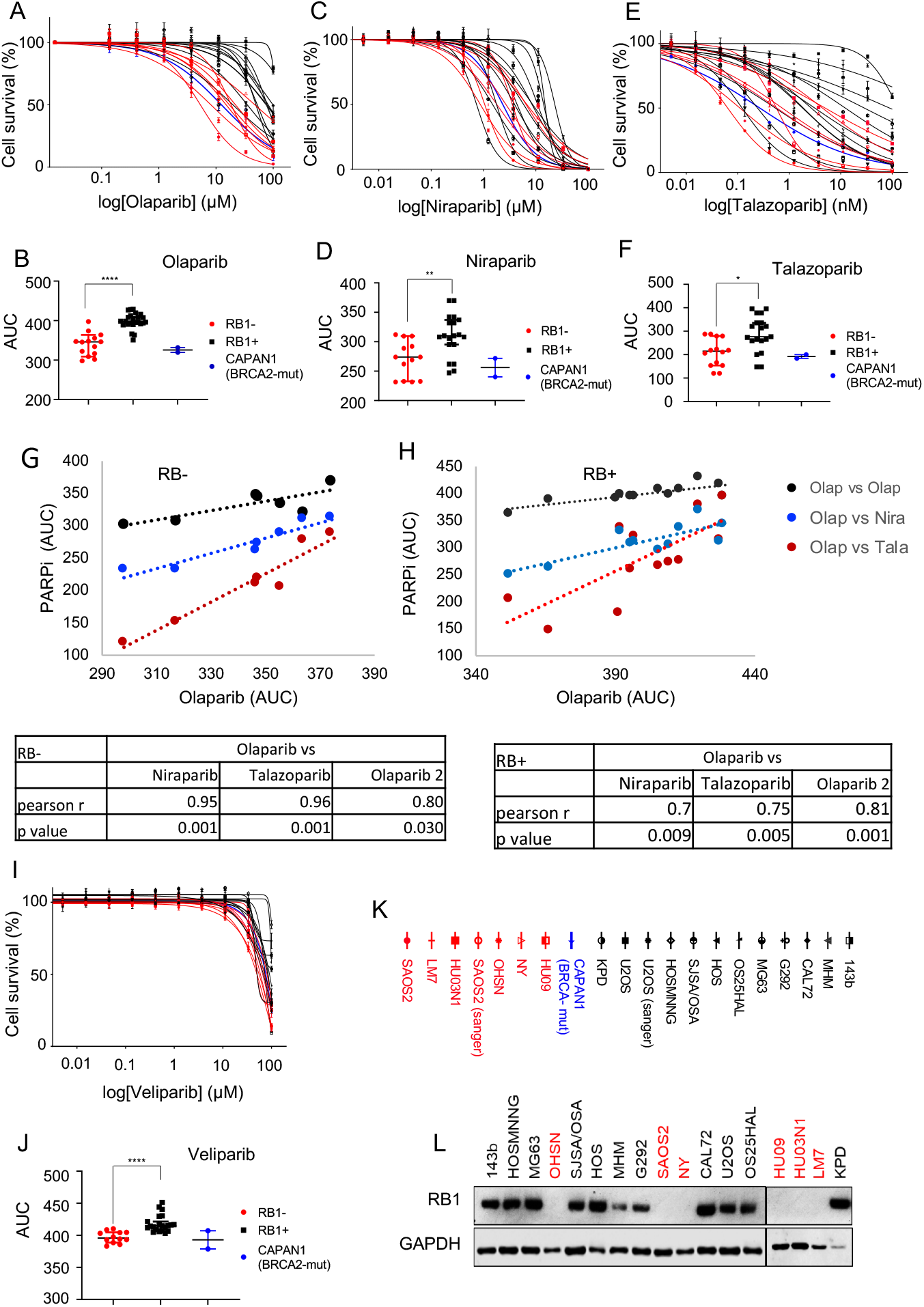
Differential PARPi sensitivities in *RB1*-mutant and *RB1*-normal osteosarcoma-derived tumour cell lines. Cell seeded in 96-well plates were treated with PARPi at concentrations indicated. Cell viability was determined 5 days of drug after exposure using resazurin reduction. **A, C, E and I)** Concentration-response curves for **A)** Olaparib, **C)** Niraparib, **E)** Talazoparib and **I)** Veliparib. Data are from one representative experiment relative to the DMSO-treated controls. Data points depict averages of three replicate samples relative to the DMSO-treated controls. Red, *RB1*-mutant and black, *RB1*-normal osteosarcoma-derived lines, blue, BRCA2-mutant pancreatic ductal carcinoma-derived CAPAN 1. **B, D, F and J)** Area under the curve (AUC) values deduced from dose response curves for individual *RB1*-mutant or *RB1*-normal osteosarcoma lines and *BRCA2*-mutant CAPAN1, treated with **B)** Olaparib, **D)** Niraparib, **F)** Talazoparib and **J)** Veliparib, depicted as scatter plots showing distribution of data for two or more individual experiments. Bars depict median and 95% confidence interval (CI), * p<0.05, **p<0.01, ****p<0.0001, using a 2-sided Mann-Whitney test. **G, H)** Pearson product moment correlation measuring the strength of a linear association between AUC data for Olaparib and AUC data for Talazoparib, Niraparib or a 2^nd^ Olaparib dataset, **G)** *RB1*-mutant osteosarcoma lines and **H)** *RB1*-normal osteosarcoma lines. Tables showing Pearson’s correlation coefficient and p values. **K)** Symbols and names for cell lines used. **L)** Immunoblotting analysis assessing the expression of RB1 in osteosarcoma-derived cell lines. GAPDH was used as loading control.

Similar results were obtained using the clinically approved but structurally unrelated PARPi niraparib (Figure 1C, D) and talazoparib (Figure 1E, F). Both yielded significantly increased median sensitivity for the *RB1*-mutant compared to the *RB1*-normal osteosarcoma group. Notably, sensitivities across the *RB1*-mutant group were greater than, or closely match<u>ed</u> that of *BRCA2*-mutated Capan-1 for all inhibitors studied (Figure 1B, D, F). High correlation coefficients and highly significant linear correlations were obtained comparing repeat assessments of the same inhibitors (Pearson r=0.92, p<0.0001 for olaparib, Pearson r =0.98, p< 0.001 for niraparib and talazoparib), (Supplementary Figure 1G-I), indicating reliability of the analysis.

Importantly, highly significant linear correlations were obtained comparing different PARPi, (Figure 1G, H), indicating that their shared activity of targeting PARP1/2 underlies the sensitivity profiles observed.

A significant association between sensitivity and *RB1*-defect was also observed using Veliparib, a PARPi that inhibits PARP1/2 catalysis but lacks the ability to trap PARP1/2 enzymes on damaged chromatin [26], [27], (Figure 1I, J), with good agreement between independent experiment (Supplementary Figure 1K). However, the differential in sensitivity was small and the dose required to affect cells viability readings high. While consistent with an increased dependency on PARP1/2 catalysis in *RB1*-mutant OS, these results indicate that PARP trapping may be an important mechanistic determinant for single agent potency, as known for *BRCA1/2* mutated cancers [22].

Clonogenic assays, scoring for the ability of cells to yield colonies in the presence of inhibitor, confirmed selective PARPi hypersensitivity in *RB1*-mutant osteosarcoma for all three PARPi (Figure 2A-B, 2D-E, 2G-H, Supplementary Table 1, raw data Supplementary Figure 2). Half maximal inhibitory concentrations (IC50) deduced from response curves revealed values in *RB1*-mutant osteosarcoma closely matching or below those determined for *BRCA2*-mutant Capan-1, with differential in median IC50 values between *RB1*-normal and *RB1*-mutant groups of 14-fold (olaparib), 5-fold (niraparib) and 8-fold (talazoparib) (Figure 2C, F, I). Superior selectivity of olaparib over niraparib has previously been observed in HRd cancers, and may relate to differences in off target activity of these different agents [28].

**Figure 2:**
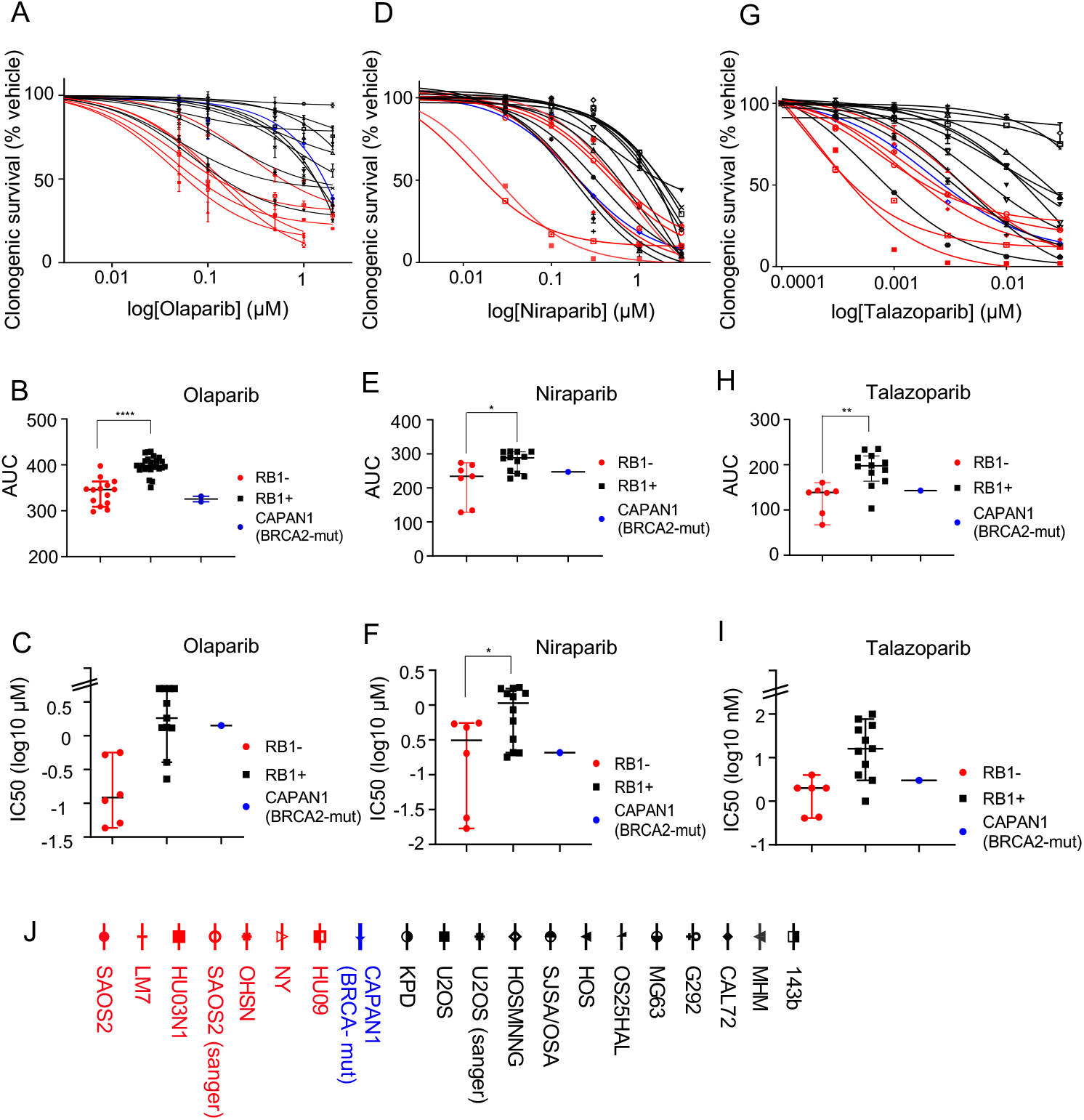
Effect of PARP inhibition on clonogenic survival. Cells were seeded into 6-well plates in the presence of vehicle (DMSO) or increasing concentrations of PARP inhibitors. Colonies arising were stained using crystal violet dye. Clonogenic survival was quantified using dye extraction. **A, D, G)** Concentration-response curves for *RB1*-mutant (red) or *RB1*-normal (black) osteosarcoma and *BRCA2*-mutant CAPAN1 (blue) after treatment with **A)** Olaparib, **D)** Niraparib and **G)** Talazoparib. Data reflect the mean +/-SD of duplicate wells for one representative experiment. **B, E, H)** AUC value comparison between *RB1*-mutant or *RB1*-normal osteosarcoma lines and *BRCA2*-mutant CAPAN1 treated with **B)** Olaparib, **E)** Niraparib, **H)** Talazoparib. AUC values are depicted as scatter plots showing distribution of data from multiple experiments. Bars depict median and 95% CI. *p<0.05, **p<0.01, ****p<0.0001 calculated using two-sided Mann-Whitney tests. **C, F, I)** Scatter plots depicting IC50 values deduced from dose response data used in B, E and H. Bars depict median and 95% CI. **J)** symbols and names for cell lines used.

Collectively the data provide evidence that *RB1* status is a predictor of single-agent PARPi sensitivity in osteosarcoma-derived cells, with sensitivity levels comparable to that of BRCA2-mutated cancer cells.

### PARPi induced cell death in *RB1*-mutant osteosarcoma

To understand how PARP inhibition acts to reduce colony outgrowth and viable cell mass in RB1-defective osteosarcoma we performed time-lapse microscopy using medium containing SYTOX death-dye, to detect cell death. Treatment with olaparib yielded a concentration-dependent increase in death-dye incorporation compared to vehicle-treated controls (Figure 3A, B) accompanied by widespread cytopathic effects (Figure 3B) in *RB1*-mutant but not *RB1*-normal osteosarcoma lines.

**Figure 3:**
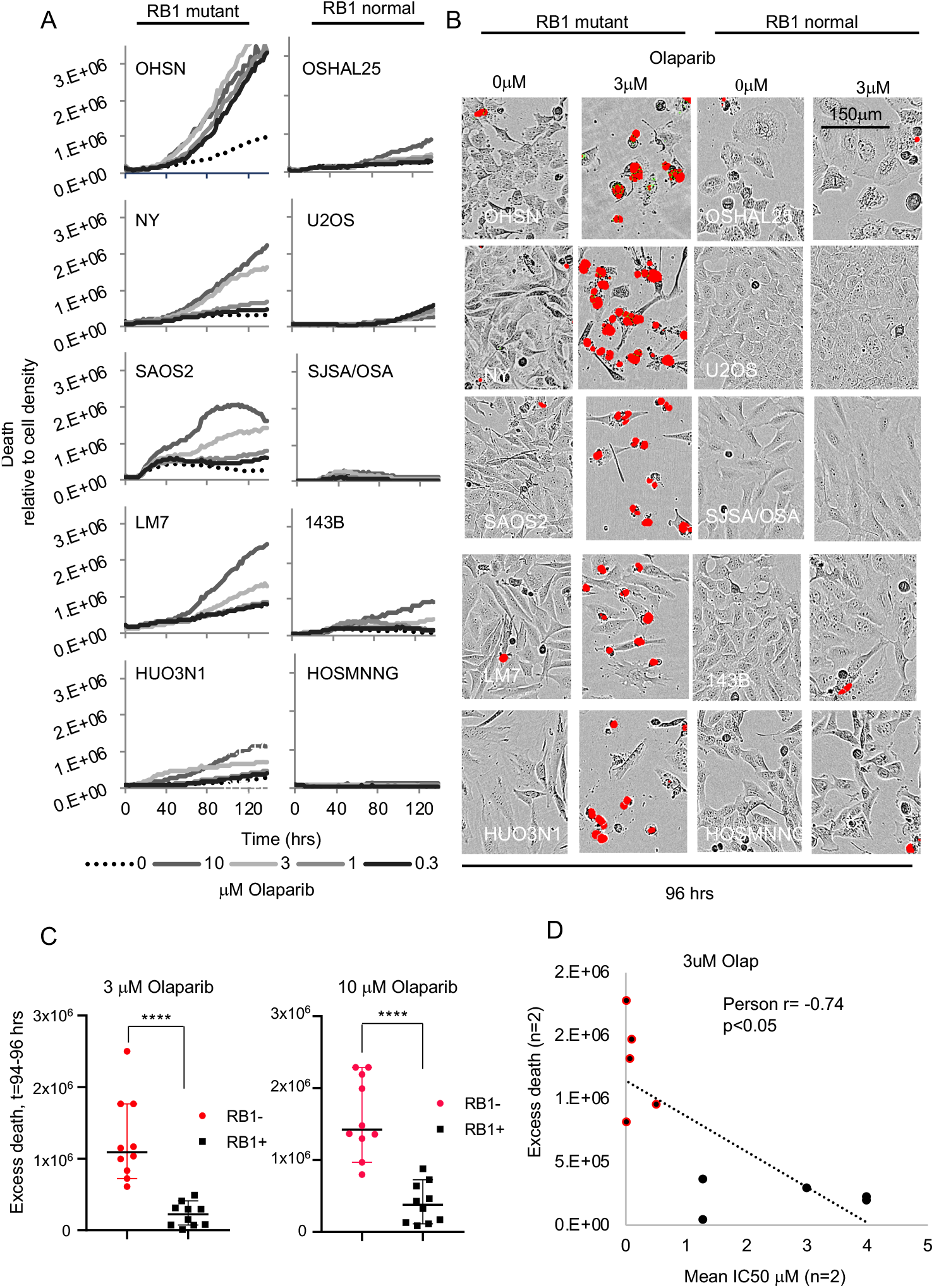
Cellular effects of PARPi treatment. *RB1*-mutant and *RB1*-normal osteosarcoma lines seeded in 96 well plates were treated with PARP inhibitor olaparib at the concentrations indicated, then subjected to time-lapse microscopy in the presence of SYTOX™ death-dye. Images were taken every two hours, recording phase contrast and death-dye fluorescence. **A)** Death-dye incorporation over time relative to cell density in *RB1*-mutant (left) and *RB1*-normal (right) osteosarcoma cancer lines. **B)** Raw images 96 hours post inhibitor addition, depicting phase contrast superimposed with death-dye fluorescence. **C)** Mean death above vehicle treated samples (excess death) at 94 to 98 hours after olaparib addition. Olaparib concentration were as indicated. **D)** Pearson product moment correlation measuring the strength of a linear association between mean excess death at 94 to 98 hours and IC50 determined using clonogenic assays for the same cell lines. Pearson’s correlation coefficient (Pearson r) and p value are indicated Data represent one exemplary experiment (A, B) or are cumulative for two independent experiments (C, D) ****p<0.0001, two-sided Mann-Whitney test.

Increased death-dye incorporation and cytopathic effects became evident between 40 and 60 hours after PARPi addition. Statistical assessment comparing death above vehicle (excess death) at 94-96 hours yielded a highly significant differential between the *RB1*-mutant and *RB1*-normal group with a strong and significant inverse correlation between death and the IC50 for the respective lines, consistent with a link between death response and antiproliferative response following olaparib treatment (Figure 3C, D).

Corroborative results were obtained using talazoparib and niraparib, confirming concentration-dependent death in *RB1*-mutant but not RB1-normal OS, with similar time to onset (40 to 60 hours) regardless of inhibitor used (Supplementary Figure 3 AC). Together these results are consistent with the enhanced sensitivity to PARPi in *RB1*-mutant osteosarcomas and identify rapid cell death as a likely key consequence of PARPi exposure in osteosarcoma cells with this genetic defect.

### PARP inhibitor sensitivity is a consequence of *RB1* deficiency

To investigate if selective PARPi sensitivity in *RB1*-mutant osteosarcomas is a consequence of RB1 loss, we depleted RB1 in the *RB1*-normal osteosarcoma line CAL72 using *RB1*-targeting small-hairpin RNAs (shRNA) (Figure 4A).

**Figure 4:**
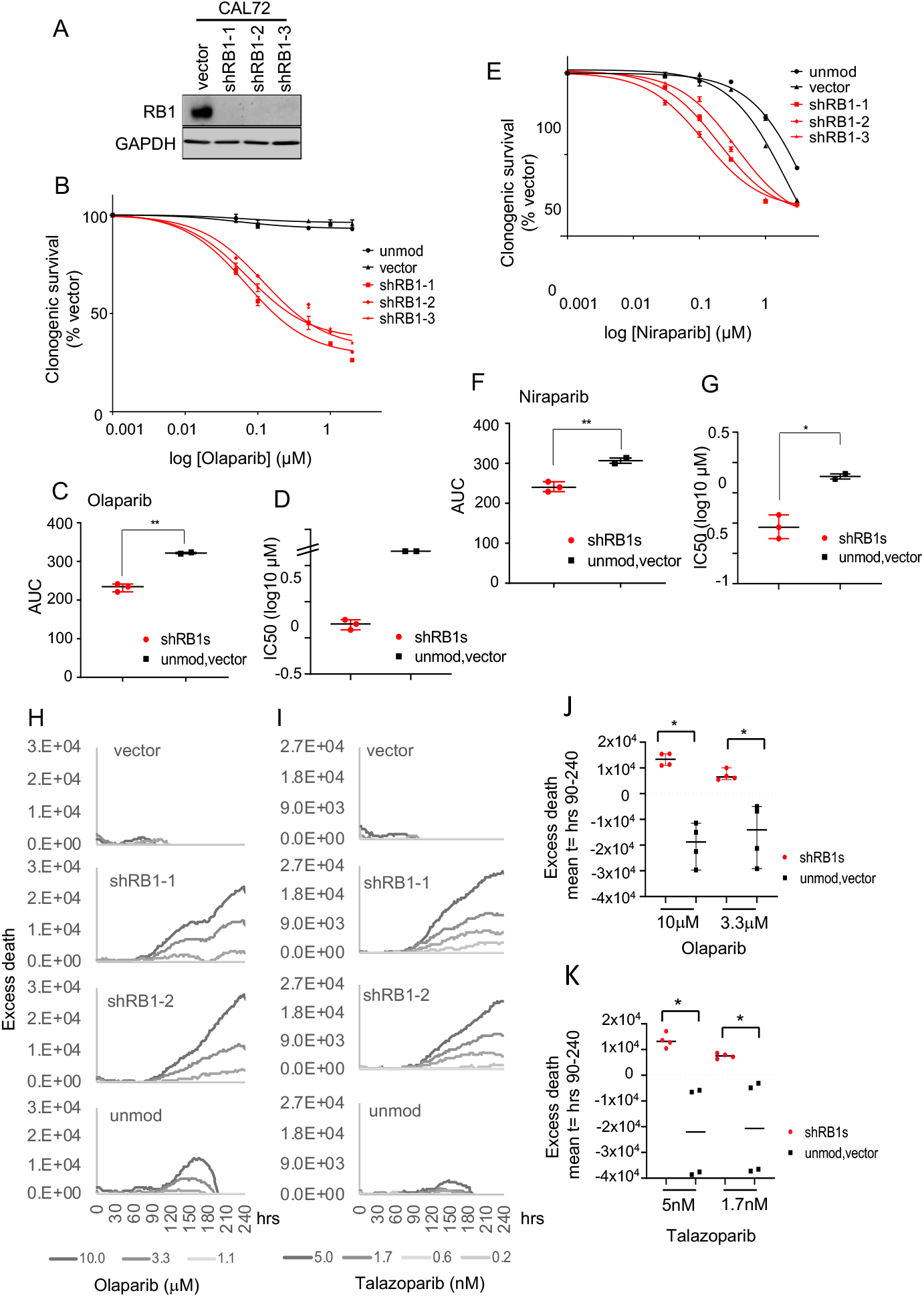
PARPi response in *RB1*-normal osteosarcoma following RB1 depletion. *RB1*-normal osteosarcoma CAL72 cells were infected with lentivirus vector encoding different *RB1*-targeting shRNAs (shRB1-1, shRB1-2 or shRB1-3) or empty vector backbone (vector) or were left unmodified (unmod.) **A)** Immunoblot analysis documenting lack of RB1 expression in CAL72 carrying *RB1-*targeting shRNAs. GAPDH was used as a loading control. **B-F)** Clonogenic survival analysis depicting concentrations-effect curves, AUC scatter plots and IC50 data for cells treated with **B, C, D)** cells treated with Olaparib or **E, F, G)** cells treated with Niraparib. Cal72 with modifications as indicated were seeded into 6 well plates and treated with PARPi using concentrations as indicated. Data reflect the mean +/-SD of duplicate wells for one representative experiment. Bars depict median and 95% CI. *p<0.05, **p<0.01, calculated using two-sided Mann-Whitney tests. **H-K)** Time-lapse microscopy assisted fate assessment. CAL72 modified as indicated were treated with PARPi and monitored for death-dye incorporation over time. **H and I)** Excess death above vehicle over time, representative raw data. **J, K**) Mean excess death at t= 90 to 240 hours. Data represent one exemplary experiment (H, I) or are cumulative for two independent experiments (J, K) *p<0.05, **p<0.01, ***p<0.001, calculated using two-sided Mann-Whitney tests. Bars depict median and 95% CI.

Clonogenic survival assays using various RB1-depleted CAL72 lines revealed a significant increase in olaparib sensitivity compared to unmodified CAL72 or controls infected with empty vector (Figure 4B, C and Supplementary Figure 4A), with IC50 values obtained in the RB1-depleted lines in the sub-micromolar range and differential in median IC50 compared to controls of >10-fold (Figure 4D and Supplementary Table 2). Consistent results were obtained in experiments using Niraparib (Figure 4E-G, Supplementary Table 2 and Supplementary Figure 4B) revealing significantly greater inhibitor sensitivity in the RB1-depleted lines, with clear, albeit smaller differential in median IC50 between groups (> 5-fold), in line with similar observations in the naturally *RB1*-mutant and *RB1*-normal osteosarcoma lines.

Notably, time-lapse microscopy revealed a significant rise in cell death in RB1-depleted CAL72 compared to control and/or maternal unmodified CAL72, that progressively increased over time and with increasing olaparib (Figure 4H, J) or talazoparib (Figure 4I, K) concentrations. Collectively, these data provide strong evidence that RB1 loss is causative and responsible for the increased hypersensitivity of *RB1*-mutant osteosarcomas to PARP inhibition.

### Mechanism of PARP inhibitor sensitivity in *RB1*-mutated osteosarcoma

Since PARPi hypersensitivity in cancers is causally linked to BRCAness/ HRd [29], we sought to determine if RB1 loss may yield BRCAness/HRd, in turn explaining the PARPi hypersensitivity observed. The inability of cells to recruit the DNA recombinase RAD51 to double-stranded DNA breaks is regarded as an indicator of BRCAness/HRd [30], [31]. We therefore assessed the ability of the *RB1*-mutant osteosarcoma lines to recruit RAD51 to DDSBs induced using ionising radiation (IR) (Figure 5A-C). To benchmark response, we included BRCA2-mutant Capan-1 defective for recruitment of RAD51 to damaged chromatin, and colorectal carcinoma HT29 cells, considered homologous recombination (HR) competent and competent for RAD51 recruitment. These experiments revealed significant DNA damage-dependent RAD51 recruitment, evidenced by an increased number of cells with >15 RAD51 foci, and a significant increase in foci numbers per cell in all the *RB1*-mutated osteosarcoma lines except for one, LM7. LM7 have previously been reported as RAD51 recruitment defective [15] thought to be linked to reduced expression of multiple HR components. As expected, inability of RAD51 recruitment was seen in Capan-1 contrasting with the substantive increase in RAD51 positive cells and significant increase in the foci numbers in HR competent HT29 following IR (Figure 5A-C).

**Figure 5:**
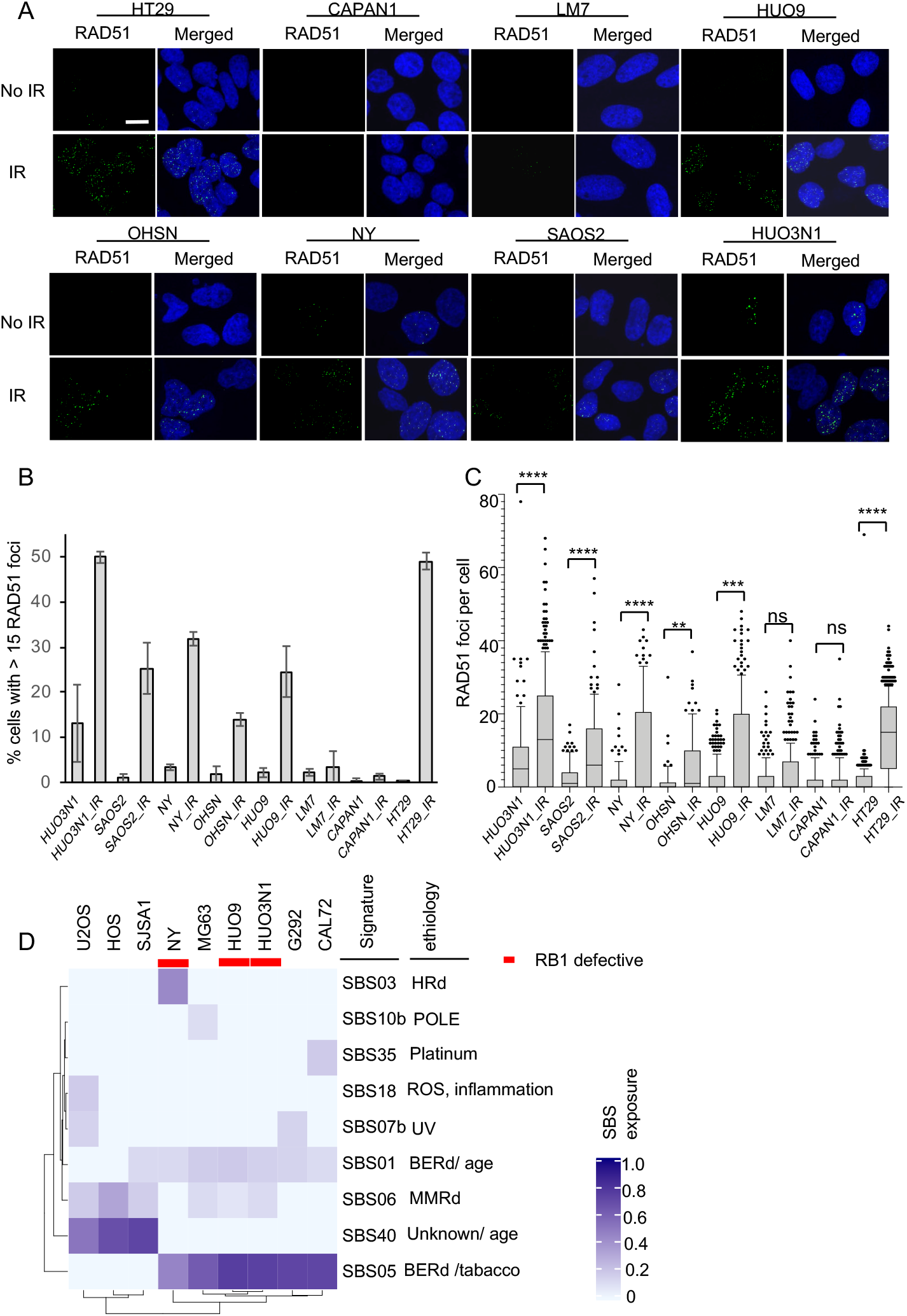
HR capability in RB1-defective osteosarcoma cell lines. **A-C)** DNA damage-dependent RAD51 recruitment in *RB1*-mutant osteosarcoma cell lines. Cells grown on glass coverslips were irradiated (4Gy) or left untreated. Cells were fixed 1 hour following IR, subjected to immunostaining for RAD51 and nuclear foci scored using confocal microscopy. A minimum of 100 cells per line were assessed across two or more experiments. **A)** Raw confocal images. Scale bar, 10 μm. RAD51 foci (green) and merged with images counter-stained for DNA using DAPI (blue). **B)** Bar chart depicting quantitation of RAD51 nuclear foci. Bars depict the % of cells with >15 nuclear foci (mean ±SEM, n>2 experiments). **C)** Box and Whiskers plot (±95%CI) depicting RAD51 foci numbers per cell. ^ns^p>0.05, **p<0.01, ***p<0.001, ****p<0.0001 using a Kruskal-Wallis test with Sidak’s multiple comparisons correction **D)** Single base substitution mutational signature analysis. Whole genome sequencing data for cell lines were downloaded from the Broad Institute’s Cancer Cell Line Encyclopedia. Signature analysis used SigProfilerMatrixGenerator.

We also performed mutation spectrum analysis (Figure 5D) using the publicly available whole genome sequence for nine of the osteosarcoma lines used. HRd in cancers is associated with a signature of somatic mutations identified as single base substitution signature 3 (SBS03), and presence of this signature provides a DNA-based measure of HRd [32]. While this analysis identified widespread presence of other signatures, evidence for exposure to HRd was only seen in one of three *RB1*-mutant lines. Notably, NY, the *RB1*-mutant line with HRd exposure, was RAD51 recruitment competent, indicative that exposure to HRd may either be historic or caused by a mechanism downstream of RAD51 recombinase recruitment. Analysis of published osteosarcoma whole exome data [33] corroborates that *RB1* defects are not significantly associated with HRd exposure (Supplementary Figure 5). HRd exposure was not detectable in 5 of 10 tumours with *RB1* mutation and had no significant linkage to *RB1* mutational status.

Together these data argue that *RB1* defects in osteosarcoma do not cause HRd/ BRCAness and hence that PARPi sensitivity in *RB1*-mutant osteosarcoma is mechanistically distinct from and not explained by outright inability to engage in HR-based DNA repair.

### Platinum sensitivity in *RB1*-mutated osteosarcoma

PARPi sensitivity in *BRCA*1/2-mutated cancer is paralleled by hypersensitivity to platinum drugs and platinum drug sensitivity is a predictor of BRCAness/HRd. Importantly, platinum drugs are an important component of clinical care in osteosarcoma. We therefore assessed if *RB1* status, that our work shows predicts PARPi sensitivity, might likewise predict platinum sensitivity in OS.

To do so we measured the sensitivity to Cisplatin across the various osteosarcoma lines using clonogenic survival assessment (Figure 6A-C, Supplementary Figure 6A, Supplementary Table 1) or day-5 viability (Supplementary Figure 6B-C). These experiments revealed similar sensitivity in *RB1*-mutant and *RB1*-normal osteosarcoma lines. Median sensitivity determined using either assay type was comparable, with no significant difference in AUC or IC50 value distributions between groups. Notably, median sensitivities closely matched that for *BRCA2*-mutated, cisplatin-hypersensitive Capan-1 [34], indicating high platinum sensitivity across osteosarcoma lines, irrespective of *RB1* status and PARPi sensitivity.

**Figure 6:**
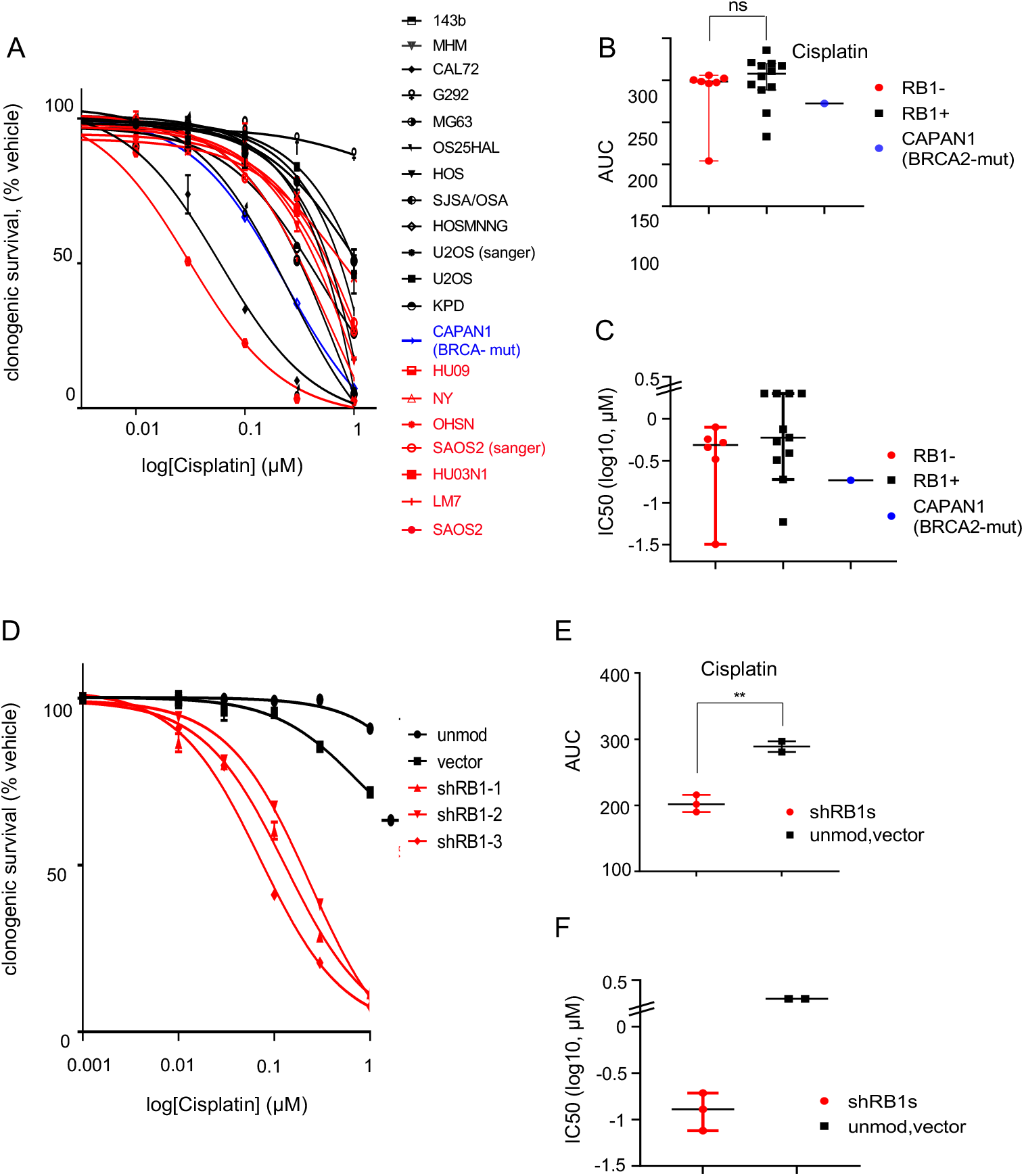
Platinum sensitivity in *RB1*-mutated osteosarcoma. Cells seeded into 6-well plates were cultured in the presence of vehicle (DMSO) or increasing concentrations of Cisplatin. Colonies arising were stained using crystal violet dye. Clonogenic survival was quantified using dye extraction. **A-C)** Platinum response in *RB1*-mutant (red) or *RB1*-normal (black) osteosarcoma and *BRCA2*-mutant CAPAN1 (blue). **A)** concentration-response curve and Scatter plots depicting **B)** AUC value comparison and **C)** log IC50 values, deduced from the concentration-response data in A). Data reflect the mean +/-SD of duplicate wells for one representative experiment. Bars in scatter plots (A, C) depict median (±95%CI), ^ns^p>0.05, calculated using two-sided Mann-Whitney test. **D-F)** *RB1*-normal osteosarcoma CAL72 cells transduced with lentivirus vector encoding different *RB1*-targeting shRNAs (shRB1-1, shRB1-2 or shRB1-3) (red) or empty vector backbone (vector) or left unmodified (unmod) (black). **D)** concentration-response curve and Scatter plots depicting **E)** AUC value comparison and **F)** log IC50 values, deduced from the concentration-response data in D). Data reflect the mean +/-SD of duplicate wells for one representative experiment. Bars in scatter plots (A, C) depict median (±95%CI), **p<0.01, calculated using two-sided Mann-Whitney test.

To assess if *RB1* defects could cause platinum sensitivity we made use of RB1-depleted CAL72. While unmodified CAL72 had modest cisplatin sensitivity (Supplementary Table 2, IC50> 1 μM), a significant and substantive sensitivity increase was seen in CAL72 in which RB1 was depleted using shRNA, based on clonogenic activity (Figure 6D-F, Supplementary Table 2, Supplementary Figure 6D) or day-5 viability (Supplementary Figure 6E, F). Hence, although platinum sensitivity is widespread amongst the established osteosarcoma lines and here not predicted by *RB1* status, these latter data argue that *RB1* defects, alike *BRCA1/2* defects, increase platinum sensitivity.

### PARPi activate DNA replication checkpoint response in *RB1*-mutant osteosarcoma

To begin to understand the causes of the PARPi hypersensitivity in *RB1*-mutant cancer cells we assessed DDSB-damage response activation in *RB1*-mutant and RB1-normal osteosarcoma cell lines. PARP inhibition prevents the ligation of single strand break and traps PARP complex on these lesions, leading to DDSB and the induction of DDSB repair signalling once cells move into S phase [35].

To assess if double strand breaks repair signalling is detectable and may selectively arise in *RB1*-mutant cells, we measured the level of the DDSB repair histone marker γH2AX using immunohistochemistry (Figure 7A, B and Supplementary Figure 7A, B).

**Figure 7:**
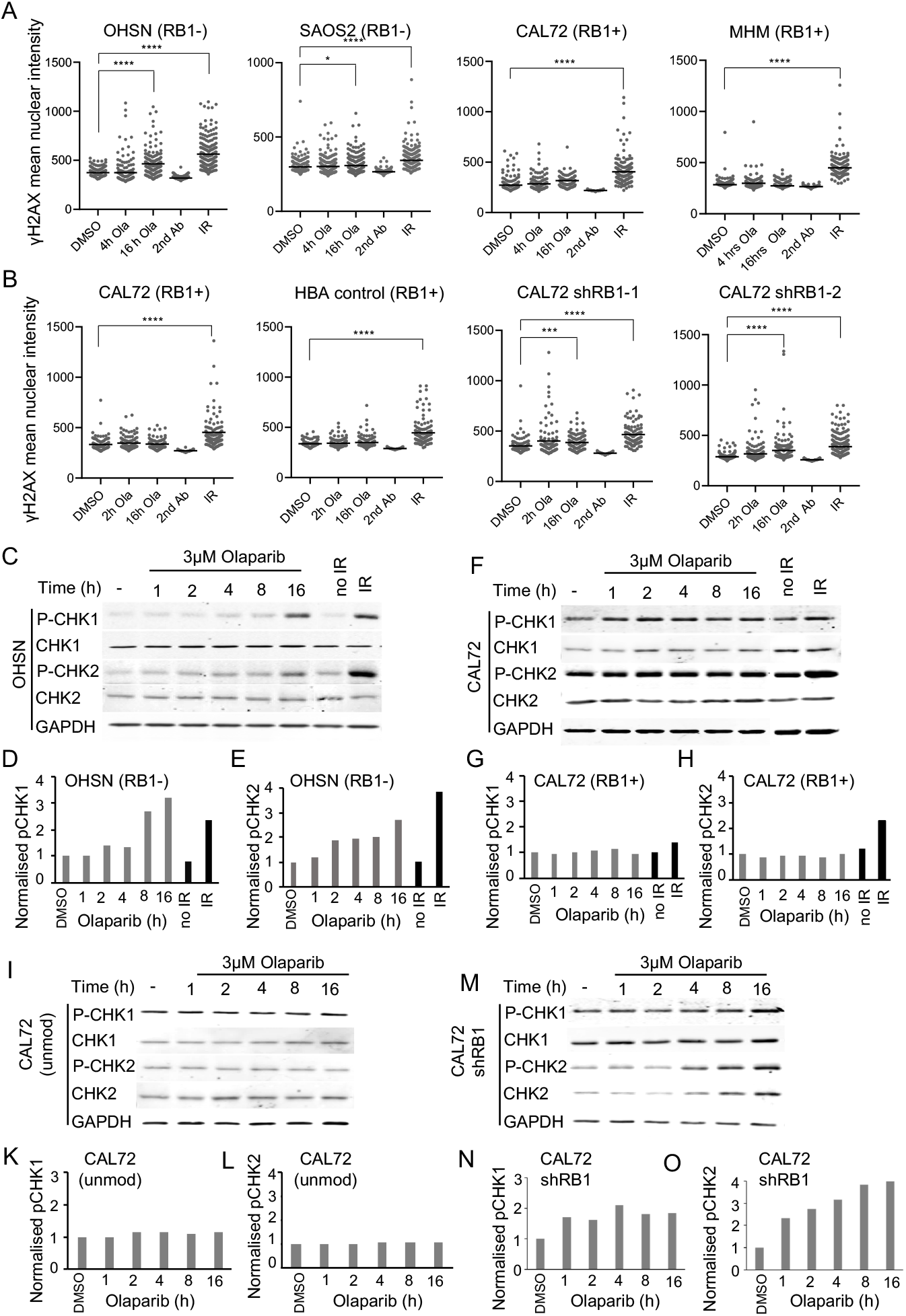
Effect of PARP inhibition on DNA damage response in cells with different RB1 status. **A-B)** DDSB repair signalling assessed using anti-phospho-Ser139-H2AX (□H2AX) immunofluorescence, **A)** in osteosarcoma cells with different RB1 status, or **B)** in *RB1*-normal osteosarcoma CAL72 transduced with lentivirus vector encoding *RB1*-targeting shRNAs shRB1-1. Cell lines after treatment with DMSO, 3μM Olaparib, or 2.5 Gy of IR. Cells were exposed to Olaparib for 2 or 4 or 16 hours or allowed to recover for 1 hour after IR. Scatter blots report distribution and mean for samples from one representative experiment, respectively. Data shown are representative for one of two or more independent experiments. **C-O)** Characterising DDSB repair checkpoint signalling assessed using immunoblot analysis assessed using phosphor(Ser345)-CHK1 (pCHK1) and phosphor (Thr68)-CHK2 (pCHK2) quantitative immunoblot analysis in **C-E)** RB1-mutant, **F-H)** *RB1*-normal osteosarcoma cells lines and **I-N)** Unmodified *RB1*-normal CAL72 or with shRNA-mediated *RB1* depletion. Cell lines after treatment with vehicle, 3μM Olaparib, or 2.5 Gy IR and exposed to olaparib for 2 or 4 or 16 hours or allowed to recover for 1 hour after IR. GAPDH was used as loading control. CHK1 and CHK2 denote immunoblot signals for pan CHK1 and CHK2. Bar graphs represent pCHK signals relative to GAPDH. Data shown are representative for one of two or more independent experiments.

We observed a robust and significant increase in γH2AX positive cells following treatment with PARPi olaparib in two different *RB1*-mutant osteosarcoma lines (Figure 7A), seen within 2 hours of treatment. Signals in cells positive for γH2AX were confined to the cell nucleus with characteristic speckled appearance, comparable in distribution and intensity to that observed following exposure to IR (Supplementary Figure 7A). No significant increase in γH2AX-positive cells compared to vehicle treatment was observed in two *RB1*-normal osteosarcoma lines albeit γH2AX-positive cells significantly increase following IR exposure (Figure 7A and Supplementary Figure 7B). Importantly, PARPi treatment induced significant γH2AX positivity following RB1 ablation in *RB1*-normal CAL72 using two differing *RB1*-targeting shRNAs (Figure 7B). No significant signal increase was seen in CAL72 that expressed irrelevant control shRNA against the alpha chain of human haemoglobin A (HBA) or were unmodified. These results indicate that canonical DDSB damage signalling ensues in response to PARP inhibition of *RB1*-mutant OS, with direct evidence that RB1 loss is a prerequisite and causative determinant in this response.

γH2AX may signify activation of distinct DNA damage response pathways, notably, ATM activated in response to DDSB, or ATR activated in response to DNA replication impairment. To delineate which of these pathways may be activated we scored for the activating modification of checkpoint kinase CHK1, selectively activated by ATR, and CHK2, linked to ATM signalling [36], using quantitative immunoblot analysis. These experiments revealed a prominent increase in CHK1 activation following PARPi treatment of the *RB1*-mutated OHSN (Figure 7C, D), which surpassed the level of activation of this kinase in response to IR in the same cells (Figure 7D). Using the same lysates, only a modest activation of CHK2 was observed, despite strong activation of CHK2 in response to IR (Figure 7C, E). PARPi treatment failed to induce significant activation of CHK1 or CHK2 in the *RB1*-normal CAL72 (Figure 7F-H), consistent with the lack of PARPi-induced γH2AX positivity in these cells. However, prominent CHK1 activation arose when RB1 was ablated using *RB1* targeting shRNA (Figure 7M-O) but not unmodified CAL72 run in parallel (Figure 7I-L). Hence PARPi treatment elicits signalling consistent with replication checkpoint activation in *RB1*-mutant cells, indicative that replication fork impairment is a key event arising in these cells.

### Requirement of DNA replication for PARP inhibitor toxicity in *RB1*-mutant cells

To address if DNA replication is a requirement for toxicity of PARP inhibition to unfold in *RB1*-mutated OS, we assessed whether preventing this process prevents PARPi-induced death in those cells. We cultured *RB1*-mutated OHSN in medium containing excess thymidine to stall replication activity during olaparib treatment (Figure 8A). Subsequently we quantified cell death measuring SYTOX dye uptake using time-lapse imaging. OHSN cells treated with olaparib whilst under thymidine-induced DNA replication block showed striking, highly significant reduction in cell death rate, compared to cells treated with olaparib in the absence of thymidine. Yet death response was restored and to levels similar to that in cycling cells when cells were released from the thymine-induced block prior to olaparib addition, (^ns^p= 0.1736) (Figure 8B-C). These results provide direct evidence that ongoing DNA replication is required for death to unfold in *RB1*-mutated osteosarcoma cells in response to PARPi treatment.

**Figure 8:**
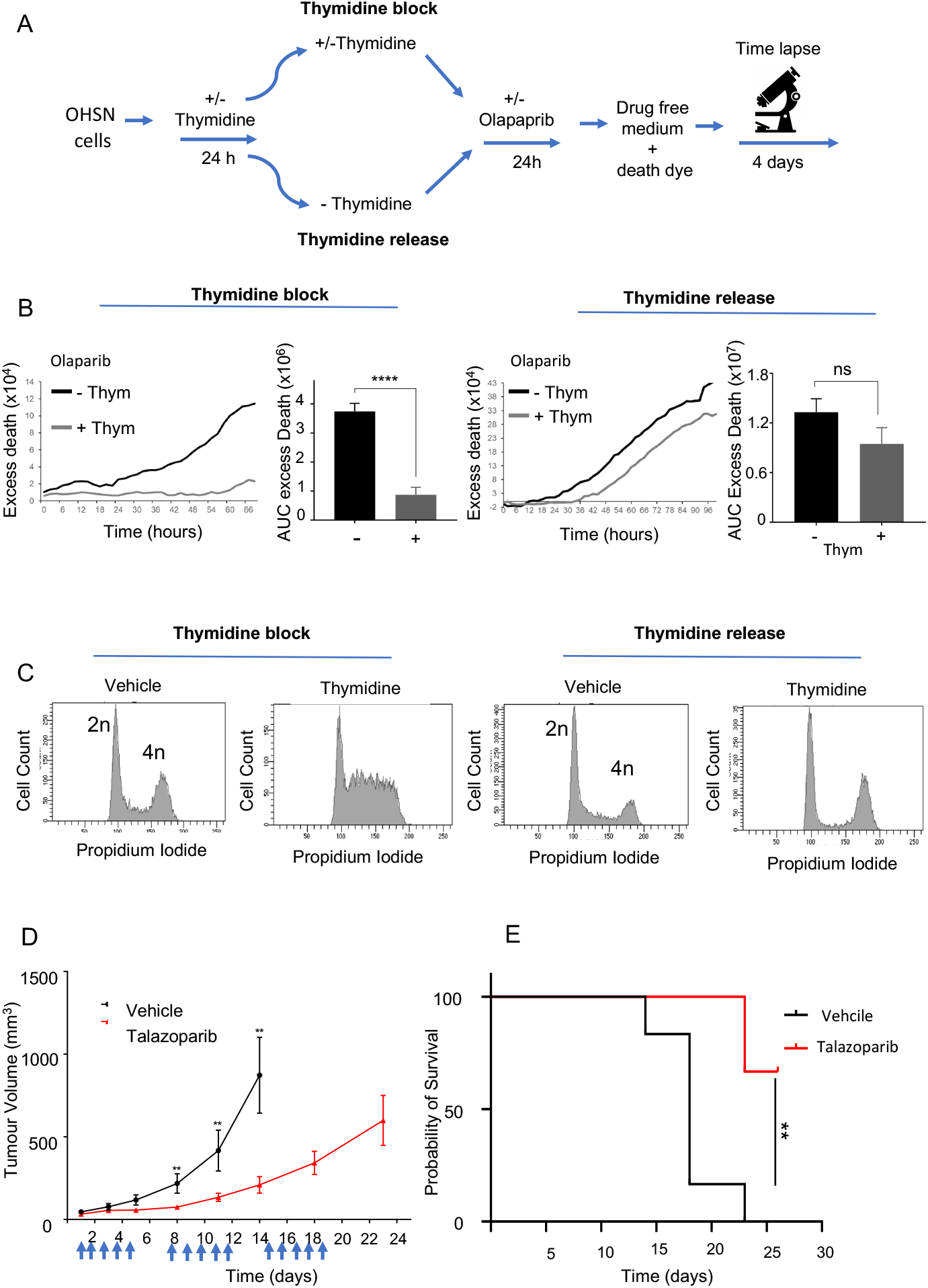
Effect of DNA replication impairment and tumour response *in vivo*. **A-C)** PARPi sensitivity following DNA replication perturbation **A)** Experiment design for assessing the role of DNA replication in PARPi sensitivity. *RB1*-mutant OHSN seeded in 96 well plates were treated as indicated, then subjected to time-lapse microscopy in the presence of SYTOX™ death-dye. **B)** Death assessed SYTOX™ death-dye incorporation. Raw traces depicting excess death above vehicle for one representative experiment, and bar graphs depicting AUC values summarizing excess death over vehicle (±SEM) for n=5 independent experiments. ^ns^p>0.05, ****p<0.0001, calculated using unpaired student *t* test. **C)** Cell cycle profiles documenting the effect of thymidine treatment on cell cycle progression. Cells seeded in parallel 6 well plates were treated as for B), then analysed using flow cytometry at 32 hour. Data shown are from one representative experiment. **D-E)** Tumour response to single agent PARPi treatment D) NRG (NOD.Cg-Rag1tm1Mom Il2rgtm1Wjl/Szj) mice carrying OHSN tumour xengrafts were treated daily (5 times per week) with the PARPi using talazoparib at 0.33 mg/kg, or empty vehicle for 3 weeks (n=6 per group). D) Tumour volumes over time (mean ±SEM, n = 6), measured at the indicated time points. Arrows indicate dosing schedule of PARPi or vehicle. **E)** Kaplan-Meier survival analysis of NRG mice bearing tumours treated with talazoparib or vehicle. **p < 0.01 calculated using a log rank (Mantel-Cox) test.

### PARP inhibitor yields robust single agent activity in *in vivo* preclinical models of *RB1*-mutant human OS

Given the substantive sensitivity of *RB1*-mutated osteosarcoma cells in cell-based experiments to PARP1/2 inhibition we sought to assess whether the observed single agent anticancer activity of PARPi extended to *in vivo* models of human OS. To this end we generated xenografts of *RB1*-mutated OHSN in immunodeficient NRG mice. Following tumour formation, mice were randomised and treated once daily for three successive 5-day periods with either vehicle (DMSO) or talazoparib at 0.33 mg/kg (Figure 8D, E).

Treatment using this schedule was tolerated with no significant impact on weight (Supplementary Figure 8) or other adverse effects. However, a highly significant reduction in tumour growth was seen in the talazoparib-treated compared to vehicle-treated mice. A significant reduction in tumour growth was apparent following the initial 5 day treatment period (**p<0.01). Importantly, while tumours in vehicle-treated mice progressed rapidly (Figure 8D), reaching the maximally allowable endpoint by 22 days, none of the tumours in talazoparib-treated mice progressed to this level within this time. Although dosing of talazoparib was discontinued on day 20 more than 70% of the tumours in talazoparib-treated mice remained within allowable limits at 26 days when observation was terminated (Figure 8E). These data provide evidence that the PAPRi sensitivity observed in cell-based assay translates into substantive single-agent anti-tumour activity yielding reduced disease progression and extended survival in mice carrying human *RB1*-mutant osteosarcoma xenografts.

## DISCUSSION

Our work identifies PARP inhibition as a synthetic vulnerability and therapeutic opportunity for RB1-mutated osteosarcoma with additional evidence that deleterious RB1 mutation may be a biomarker of clinically relevant PARP1/2 inhibitor sensitivity in other cancers. PARPi are in current clinical use, with notable effect on quality of life and overall survival in multiple cancers [37]. Their existing clinical utility highlight the imminent opportunity for clinical translation footing of the finding we here report.

Patient selection in current clinical applications relies on evidence of HRd in cancer tissue [38], [39]. However genomic or functional evidence for frank HRd was not detectable or significantly associated with RB1 loss in cancer, which would have precluded selection of these cancers from current PARPi treatment regiments.

Our work documents that enforced RB1 loss causes clinically meaningful sensitivity (i.e. sensitivity akin to that seen in *BRCA1/2*-defective Capan-1) in an otherwise PARPi insensitive osteosarcoma line, providing proof of concept for a direct contribution of RB1 loss to the selective PARPi sensitivity observed. The lack of frank HRd in cancer cell lines with RB1 loss raises questions as to the mechanism that underlies their sensitivity. Our work positively identifies PARPi trapping and active DNA replication as mechanistic prerequisites for sensitivity, paralleling observations in cancers with HRd [26], [40]. These observed similarities argue for a shared inability in cancers with HRd or RB1 loss to avert the lethal consequence of replication fork collapse, caused by trapped PARP complex and known to underly PARPi inhibitor sensitivity caused by HRd.

Published data propose a role of RB1 in HR, entailing E2F1-dependent recruitment of chromatin remodelling activity to sites of DNA damage [18], albeit, the scale of HRd arising through this mechanism has not been assessed. It is conceivable that localised HRd arising within subgenomic contexts arises, and although not detected in genome wide mutation spectrum analysis could cause a synthetic lethality interaction between RB1 loss and PARP inhibition. Other evidence links chromatin processes, including defective DNA cohesion and chromatin remodelling to PARPi sensitivity [41], [42]. Defects in these processes are known to result from RB1 loss [43], which in turn could explain the observed sensitivity phenotype.

While our work advocates the use of PARPi in *RB1*-mutated osteosarcoma, comprehensive preclinical validation, including how PARPi should be best integrated into the current management of osteosarcoma, will be of likely paramount importance to ensure clinical benefit.

PARPi are rapidly moving into first line clinical use in patients with HRd ovarian cancers and considerable efforts are underway to extend their use to other cancers. Most pertinent to the work reported here is the planned assessment of PARPi within the paediatric MATCH study (NCT03233204), a large scale precision medicine trial in children, adolescents, and young adults with advanced cancers including osteosarcoma, with use of BRCA1/2 mutation or HRd for patient selection. The highly penetrant hypersensitivity in RB1-mutant osteosarcoma cells shown here, combined with the currently limited options in patient with such tumours advocates expansion of such assessment to include RB1-mutated disease.

## MATERIAL AND METHODS

### Cell lines, Chemicals and Antibodies

The osteosarcoma tumour cell lines were described previously [15]. PARPi and cisplatin were purchased from Selleck Chemicals. Antibodies and shRNAs are summarised in supplementary materials. Mutation spectrum analysis was as described [32].

### Assessment of drug response

Drug sensitivity was assessed in 96-well plates based on Resazurin reduction 5 days following drug addition. Clonogenic survival assays and immunofluorescence staining were performed as described [44]. Survival data were plotted using a three-parameter regression curve fit in GraphPad Prism 8 software. Time-lapse microscopy was performed in 96-well plates as described in [45] using an IncuCyte ZOOM live cell analysis system (Essen Bioscience). For cell cycle analysis, cells were fixed in 70% ethanol, stained using propidium iodide and analysed using flow cytometry. Immunoblot analysis used whole-cell protein extracts prepared by lysis of cells into 0.1% SDS, 50 mM TRIS-HCL, pH 6.8, containing protease and phosphatase inhibitors (ThermoFisher Scientific, UK). *In vivo* experiments were carried out under UK Home Office regulations in accordance with the Animals (Scientific Procedures) Act 1986 and according to United Kingdom Coordinating Committee on Cancer Research guidelines for animal experimentation [46] with Animal Welfare Ethical Review Body (AWERB) approval. Tumour-bearing NGS mice were randomly assigned to treatment once tumours reached 100 mm^3^. Tumour growth was assessed twice weekly using digital callipers. Assessments were terminated in accordance with AWERB guidelines.

### Statistical Analysis

Statistical hypothesis testing was performed using Microsoft Excel or GraphPad Prism. Statistical tests used are named within the text. Differences with p<0.05 were considered statistically significant.

Further detailed information on all methods is provided in the Supplementary Materials (available online).

## Authors contribution

GZ, NP, SS and SM contributed to the design of experiments; SM lead the project; GZ and SM wrote the manuscript; GZ, CAM, CM, RMA, MD, and CM performed specific components of the experimental work and analyzed associated data; CS performed the bioinformatics-based mutation signature analysis; J.M. assisted with the microscopy analysis.

## ACKNOWLEDGEMENT

The work was supported by grants from Children with Cancer. GZ and CAM and MD were supported by Children with Cancer UK (ref.17-244), with additional support through philanthropic giving by charities aid foundation account A80105053. CM was supported through a Cancer Research UK studentship (ref.C416/A23233). The funders had no role in design of the study, the collection, analysis, and interpretation of the data, the writing of the manuscript, or the decision to submit the manuscript for publication.

## SUPPLEMENTARY FIGURES AND LEGENDS

**Supplementary Figure 1:**
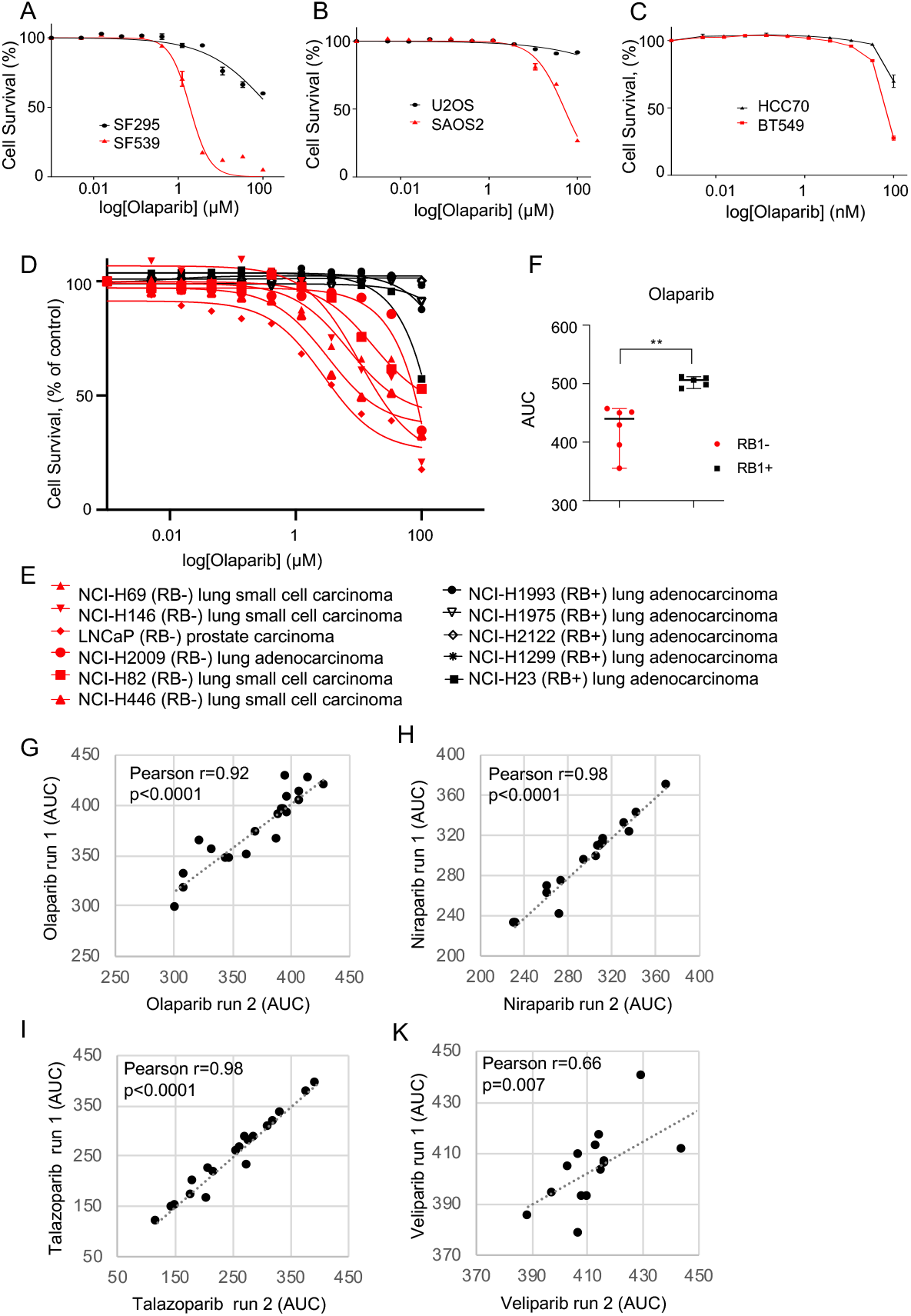
Differential PARP inhibitor (PARPi) sensitivities in RB1-mutant and RB1-normal tumour cell lines. Cell seeded in 96-well plates were treated with PARPi at concentrations indicated. Cell viability was determined 5 days of drug after exposure using resazurin reduction. **A-C)** Histiotype-matched cancer cell lines pairs with differing *RB1* mutation status were assessed for sensitivity to PARPi olaparib. Concentration-response curves were generated by calculating the decrease in resazurin signal in PARPi-treated samples relative to the DMSO-treated controls for **A)** glioblastoma, **B)** osteosarcoma, **C)** breast. Data points depict averages of three replicate samples for one of two or more independent experiments. Red, *RB1*-mutant and black, *RB1*-normal status. **D-F)** Poly-cancer cell line panel, with **D)** concentration-response curve, **E)** Symbols, names and histiotype of cell lines used for D), F) Scatter blot for AUC values deduced from D), comparing between RB1-mutant and RB1-normal tumour cell lines. Data points represent averages of three replicate samples for one of two independent experiments. Error bars indicate median and 95% CI. Red, *RB1*-mutant and black, *RB1*-normal status. ** p<0.01 calculated using a two-sided Mann-Whitney test. **G-J)** Pearson product moment correlation measuring the strength of a linear association between AUC values deduced from day-5 viability concentration response curves determining for osteosarcoma-derived cell lines shown in Figure 1. Data compare two independent experiments involving treatment, with **G)** olaparib, **H)** niraparib, **I)** talazoparib and **K)** veliparib. Pearson’s correlation coefficient r, and p-values relating to r, are shown. Data related to Figure 1

**Supplementary Figure 2:**
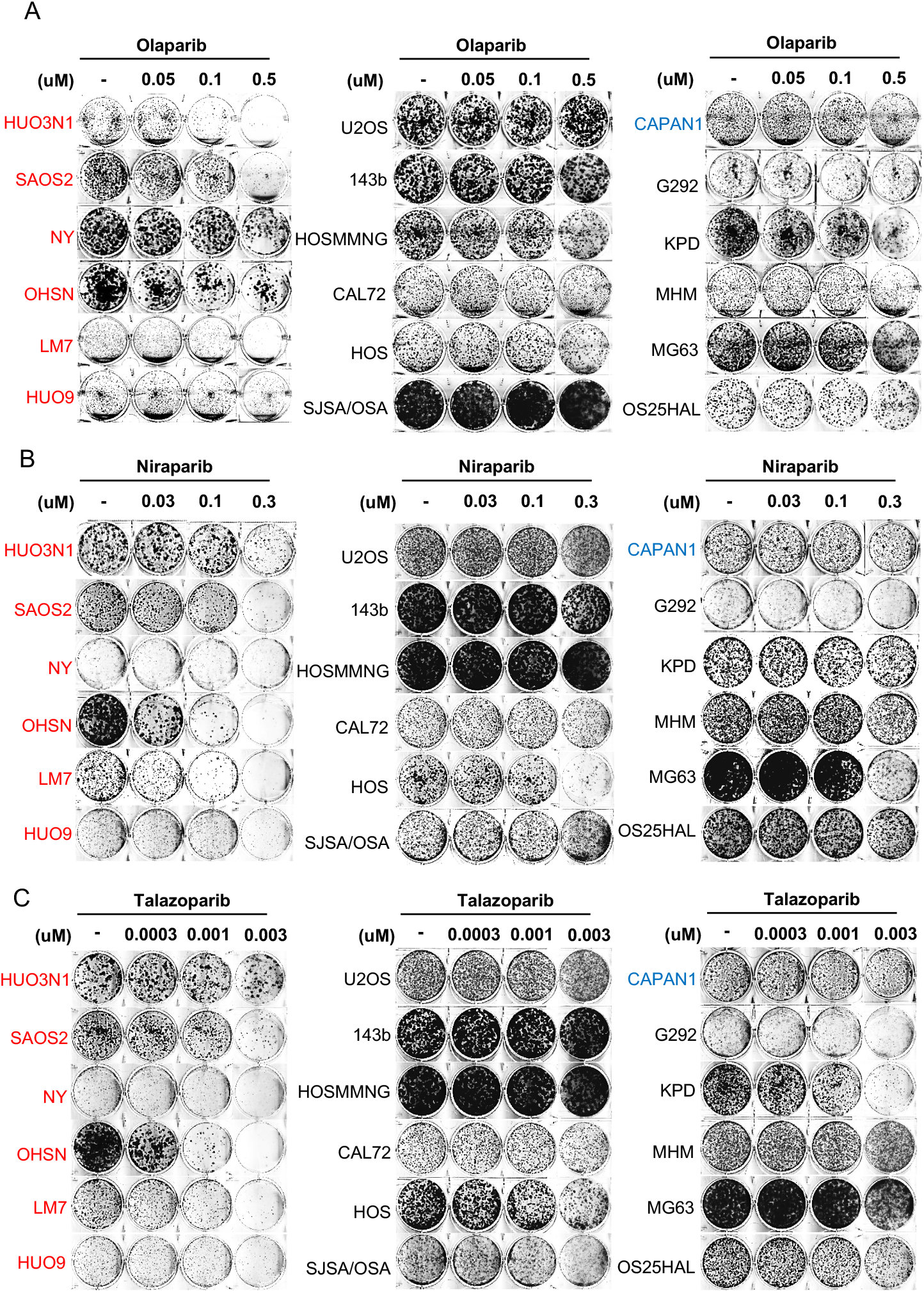
Effect of PARP inhibition assessed using clonogenic survival assays and depicting raw plate images. Osteosarcoma tumour cell lines differencing in RB1-status were seeded into 6-well plates in the presence of vehicle (DMSO) or increasing concentrations of three PARP inhibitors. Cells were treated in duplicate, fixed after 12-14 days and stained using crystal violet dye. Representative raw plate images of cell colonies for osteosarcoma lines stained with crystal violet after treatment with **A)** olaparib, **B)** niraparib and **C)** talazoparib. Red, *RB1*-mutant and black, *RB1*-normal status. Data related to Figure 2

**Supplementary Figure 3:**
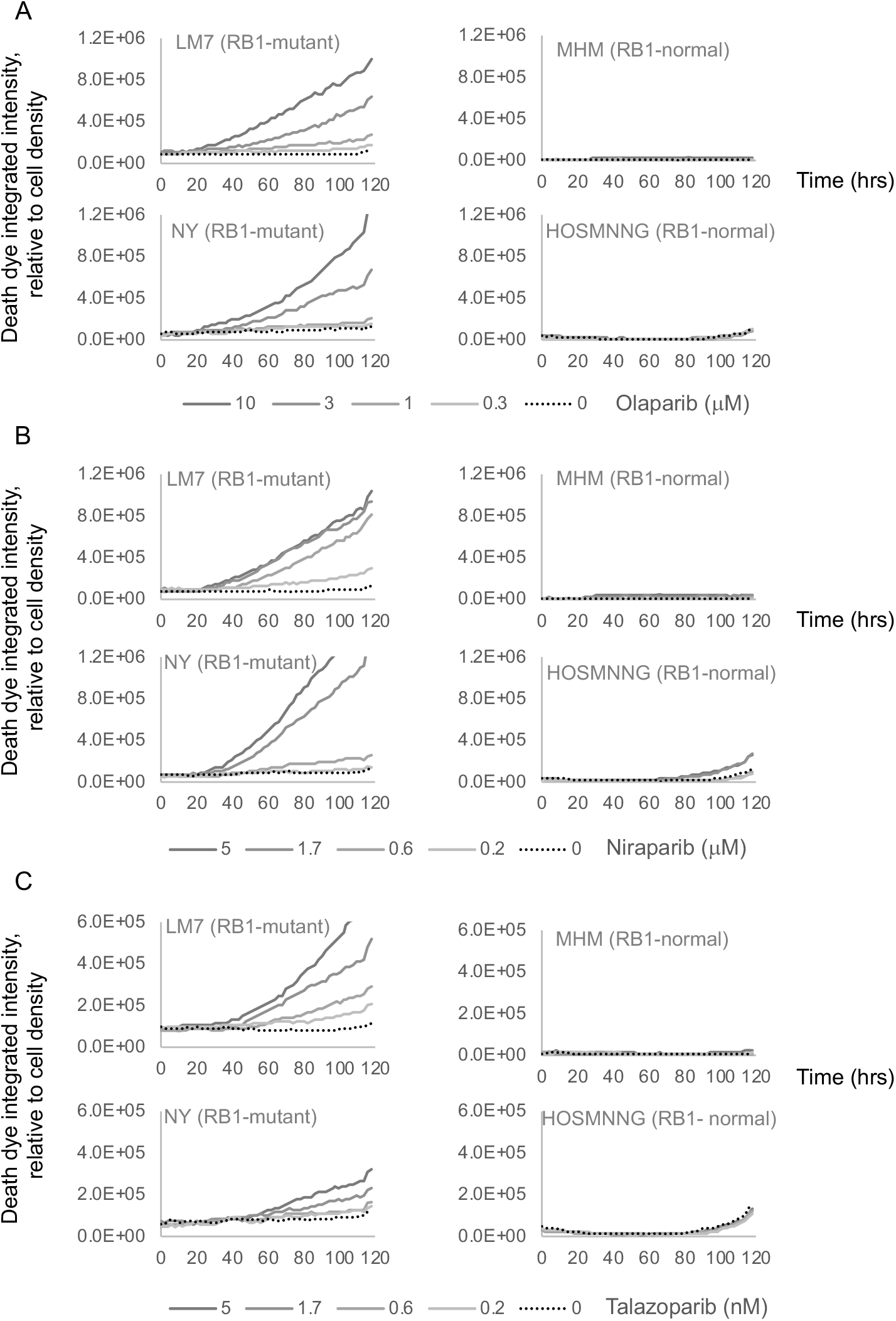
Cellular effects of PARPi treatment. RB1-mutant (LM7 and NY) and RB1 normal (MHM and HOSMNNG) osteosarcoma cell lines were treated in parallel with PARP inhibitors olaparib, niraparib and talazoparib, then subjected to (time-lapse microscopy in the presence of SYTOX™ death-dye. Images were taken every two hours, recording phase contrast and death-dye fluorescence. Inhibitors wene need at an equipotent dose range established using clonogenic survival. **A-C)** Graphs depicting death-dye incorporation over time relative to cell density with **A)** cells treated with olaparib, **B)** cells treated with niraparib, **C)** cells (treated with talnzoparib. Data represent one exemplary experiment of two or more independent assessments.

**Supplementary Figure 4:**
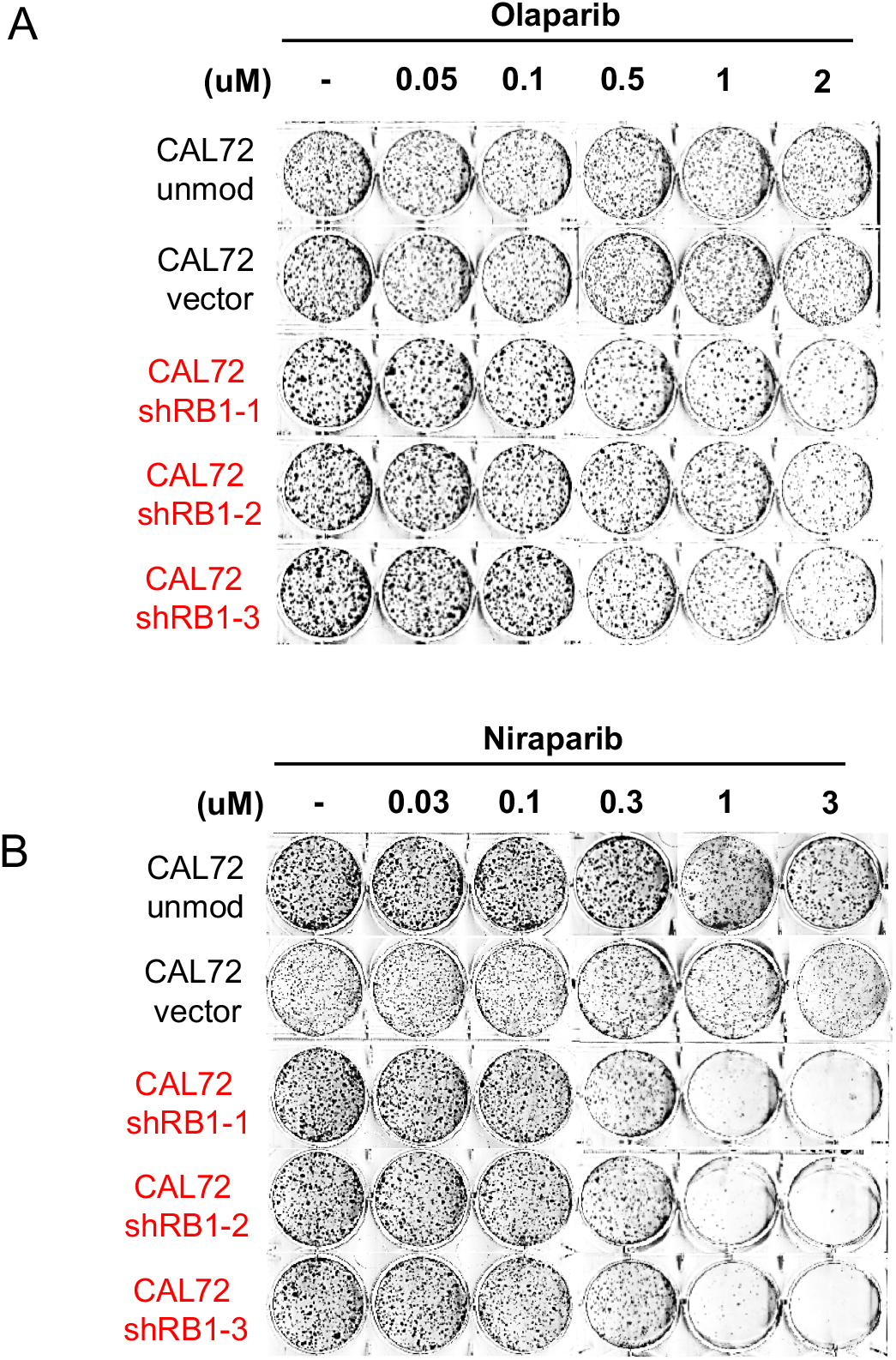
Effect of PARP inhibition in *RB1*-normal osteosarcoma CAL72 following RB1 depletion. Representative raw plate images illustrating clonogenic survival in *RB1*-normal osteosarcoma CAL72 cells with ShRNA-mediated *RB1* ablation. Cells were treated with **A)** olaparib, **B)** niraparib. CAL72 cells transduced with lentivirus vector encoding *RB1*-targeting ShRNAs shRB1-1, shRB1-2 or shRB1-3, or empty vector backbone (vector) or left unmodified (unmod).data related to Figure 4D-F.

**Supplementary Figure 5:**
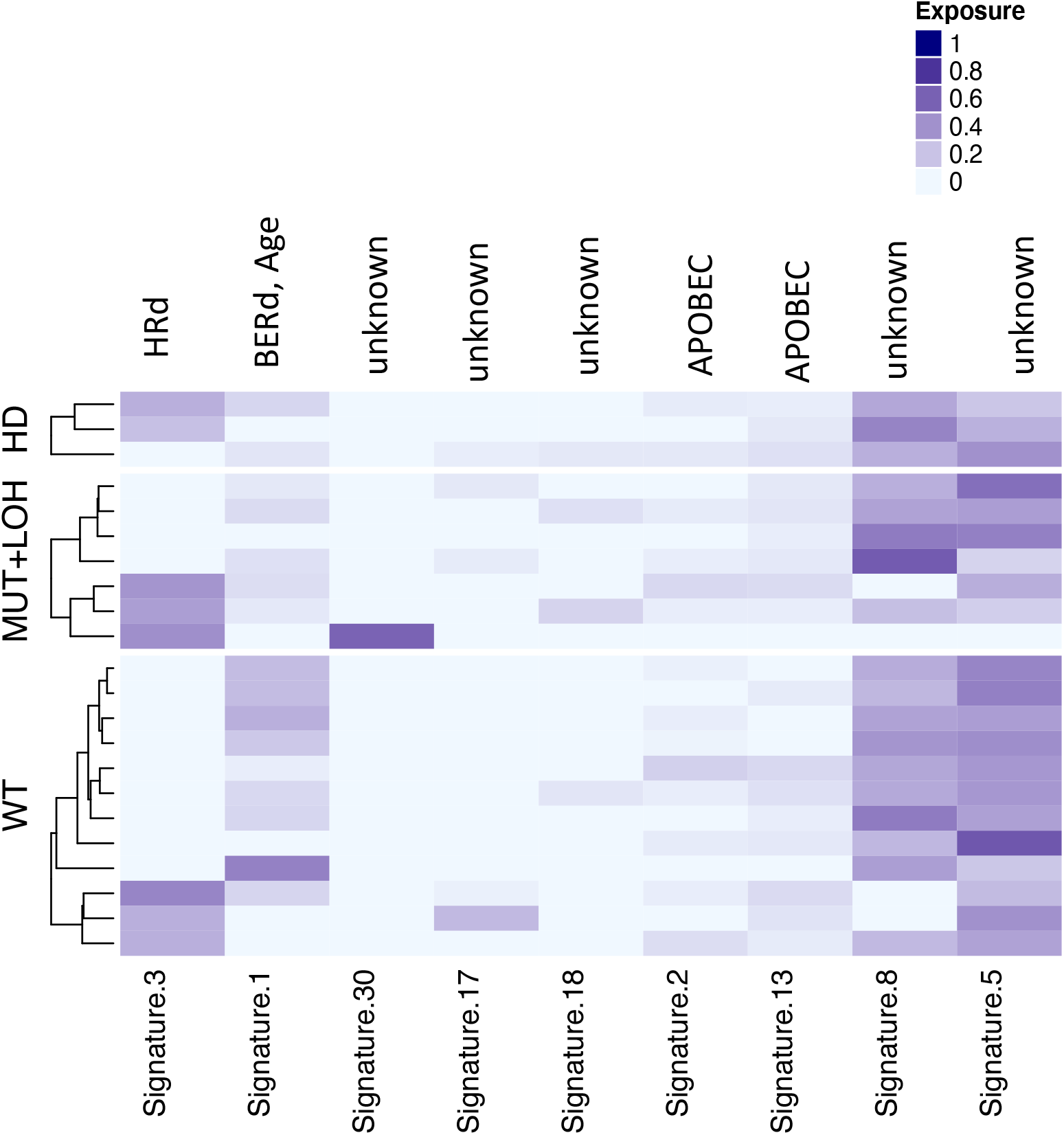
Mutational signature analysis for published osteosarcoma whole exome data. Mutational data from 37 previously published whole genome sequenced osteosarcomas was downloaded data {Behjati, 2017 #29}. Cosmic V2 signature classification with associated etiology prediction is shown. Etiology terms; Homologous recombination (HRd), Base excision repair defect (BERd), apolipoprotein B mRNA editing enzyme (APOBEC). Homozygous deletions, CN= {0,0}, and loss-of-heterozygosity events, CN= {>0,0} and mutational data were called at *RB1* to determine RB1 mutation status.

**Supplementary Figure 6:**
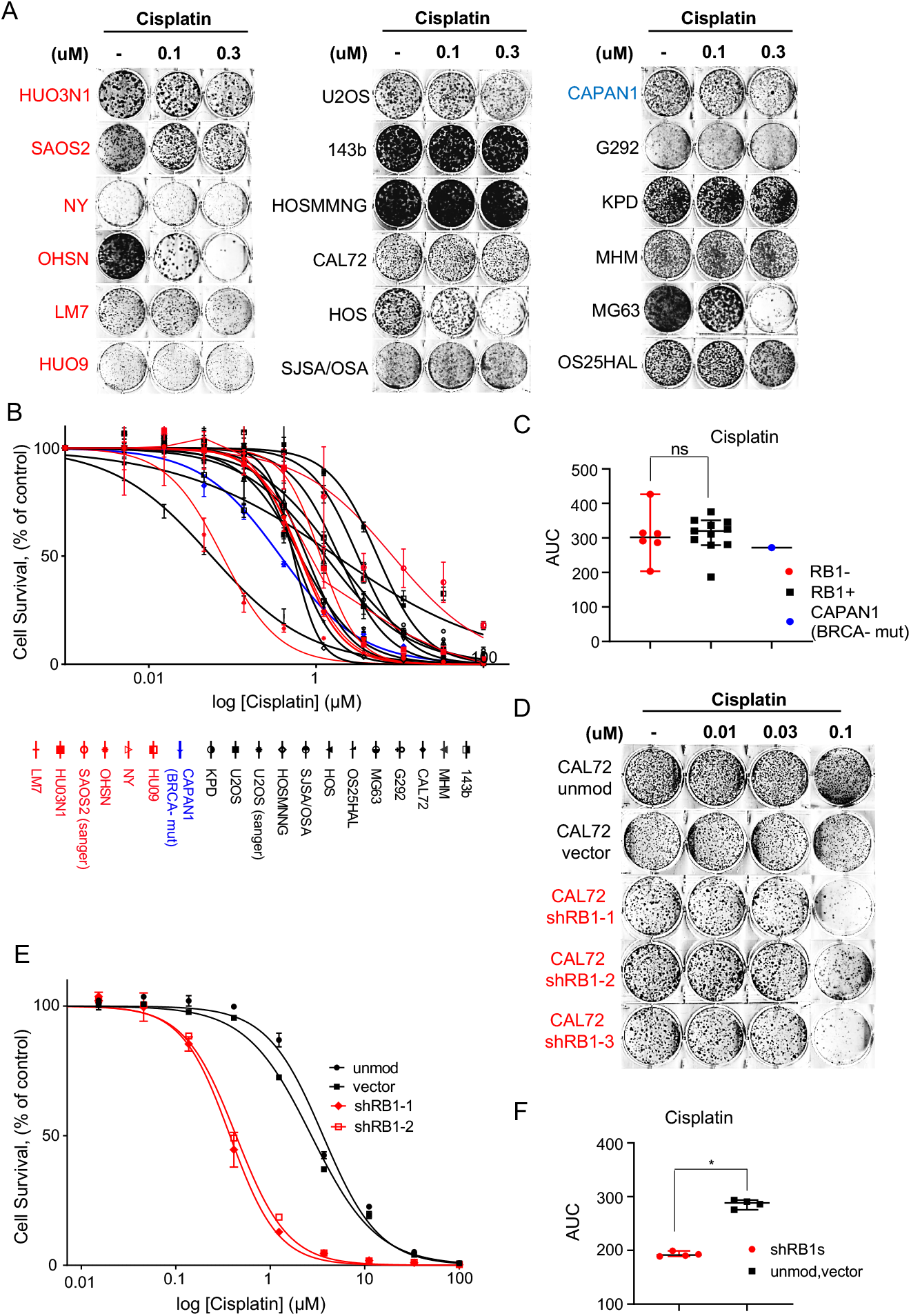
Platinum sensitivity in *RB1*-mutated osteosarcoma. **A-C)** Platinum response in *RB1*-mutant (red) or *RB1*-normal (black) osteosarcoma lines. **A)** Representative raw plate images illustrating clonogenic survival following culture in the presence of increasing concentrations of Cisplatin. Data related to Figure 6A-C. **B-C)** day-5 survival, determined using resazurin reduction. Graphs depict **B)** Concentration-response curves, reflecting mean +/-SD of triplicate wells for one representative experiment and **C)** AUC value comparison. Bars depict median (±95%CI), ^ns^p>0.05, calculated using two-sided Mann-Whitney test. Data are representative of two or more independent experiments. **D-F)** Platinum response in *RB1*-normal osteosarcoma CAL72 cells with ShRNA mediated *RB1* ablation. **D)** Representative raw plate images illustrating clonogenic survival following culture in the presence of increasing concentrations of Cisplatin. Data related to Figure 6D-F. **E-F)** day-5 survival in *RB1*-normal osteosarcoma CAL72 cells with shRNA mediated *RB1* ablation, determined using resazurin reduction. Graphs depict**, E)** Concentration-response curves, reflecting mean +/-SD of triplicate wells for one representative experiment, and **F)** AUC value comparison. Bars depict median (±95%CI), *p<0.05, calculated using two-sided Mann-Whitney test. Data are representative of two or more independent experiments.

**Supplementary Figure 7:**
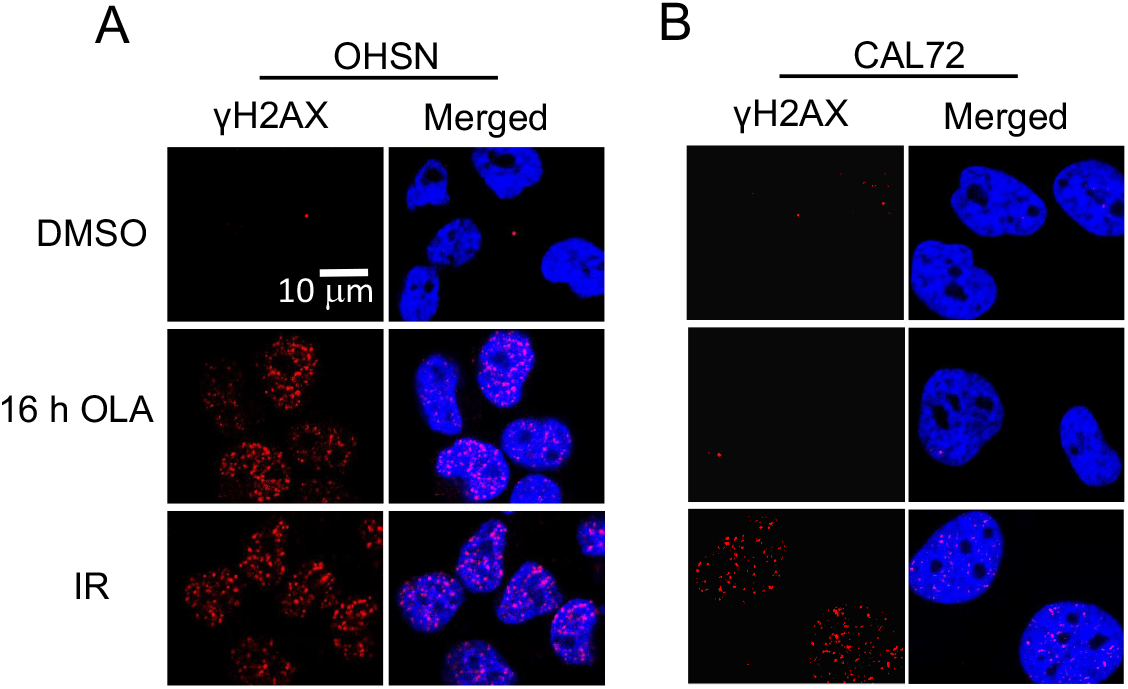
Effect of PARP inhibition on DNA damage response in cells with different RB1 status. **A-B)** Representative images for anti-γH2AX immunostaining. **A)** RB1-mutant OHSN and **B)** RB1-normal CAL72 OS. Cells were either irradiated (2Gy) and fixed 1 hour later, or treated with 3 μM of Olaparib or vehicle (DMSO) for 16 hrs. Cells were immunostained with γH2AX antibodies. Cell nuclei were counterstained with DAPI. Scale bar, 10 μm. Data related to Figure 7A, B.

**Supplementary Figure 8:**
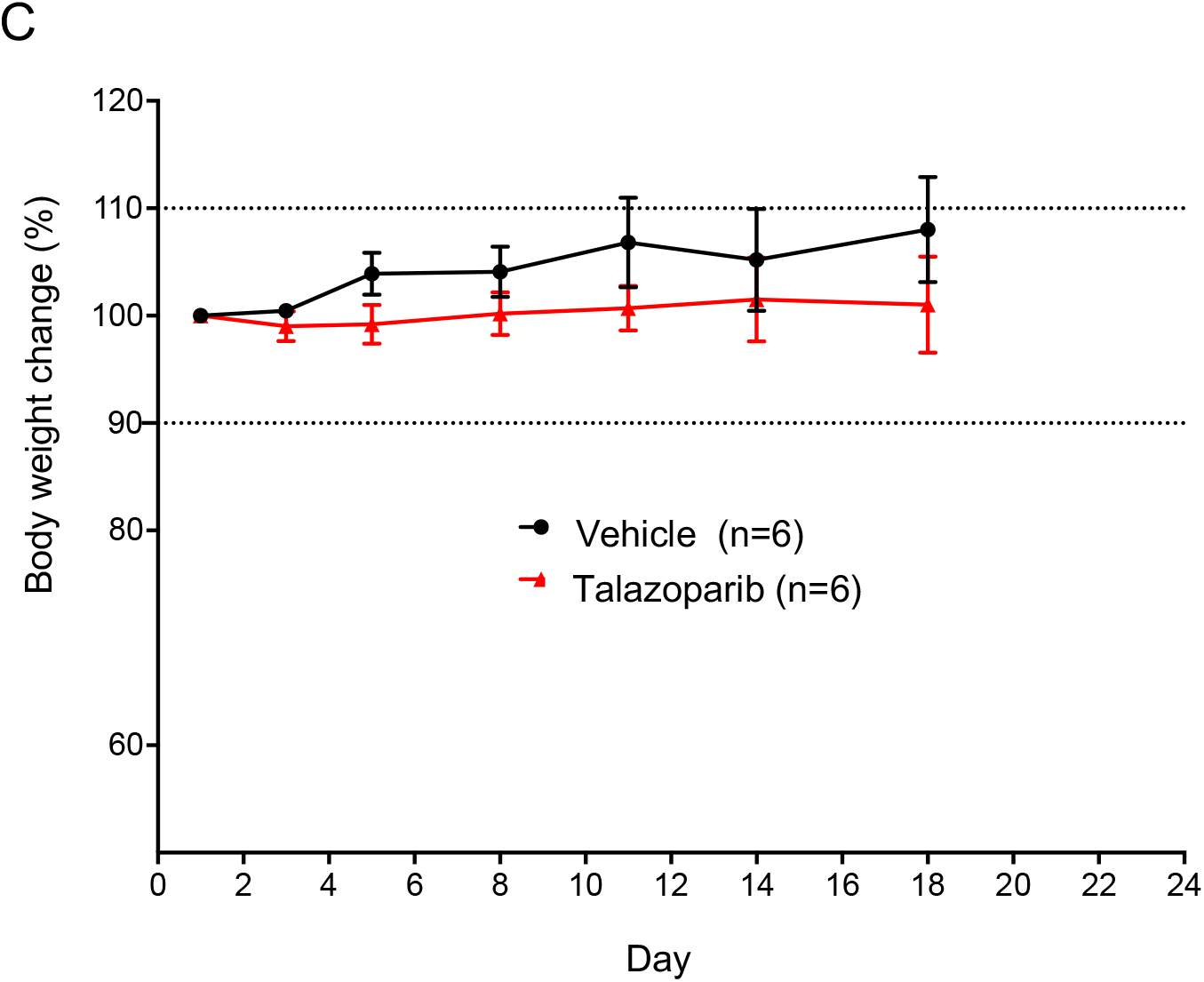
Effect of PARPi treatment on OHSN xenograft-bearing mice. Body weight over time in mice treated 0.33 mg/kg talazoparib or vehicle p.o. (mean ±SD, n = 6 per group). Data represent % change in weight relative to treatment start, lines mark 10% differential boundaries. Data related to Figure 8D, E.

## SUPPLEMENTAL TABLES

**Supplementary Table 1:** Inhibitory constant 50 (IC50) values based on inhibition of clonogenic activity. Cell lines grouped by genetic status (RB-) RB1-mutant, (RB+) *RB1*-normal, (BRCA) *BRCA2*-mutated. Median denotes median IC50 values calculated for the respective groups. n.d.= not determined. § denotes cases where an IC50 could not be extrapolated from the concentration response data obtained.

| Cell lines | Olaparib | Cell lines | Niraparib | Cell lines | Talazoparib | Cell lines | Cisplatin |
| --- | --- | --- | --- | --- | --- | --- | --- |
|  | IC50 (μM) |  | IC50 (μM) |  | IC50 (nM) |  | IC50 (μM) |
| (RB1-) |  | (RB1-) |  | (RB1-) |  | (RB1-) |  |
| HUO3N1 | 0.04 ± 0.02 | LM7 | 0.02 | OHSN | 0.41 | OHSN | 0.03 |
| OHSN | 0.05 ± 0.03 | OHSN | 0.02 | LM7 | 0.43 | HUO3N1 | 0.33 |
| LM7 | 0.11 ± 0.0q | SAOS2 | 0.20 | SAOS2 | 2.00 | NY | 0.46 |
| NY | 0.13 ± 0.01 | NY | 0.48 | NY | 2.00 | LM7 | 0.52 |
| SAOS2 | 0.52 ± 0.53 | HUO3N1 | 0.54 | HUO9 | 2.00 | SAOS2 | 0.57 |
| HUO9 | 0.56 ± 0.03 | HUO9 | 0.55 | HUO3N1 | 4.00 | HUO9 | 0.79 |
| Median | 0.12 | Median | 0.34 | Median | 2.00 | Median | 0.49 |
| (RB1+) |  | (RB1+) |  | (RB1+) |  | (RB1+) |  |
| G292 | 0.23 ± 0.12 | MG63 | 0.18 | KPD | 1.00 | HOS | 0.06 |
| KPD | 0.40 ± 0.02 | HOS | 0.21 | HOS | 3.00 | MG63 | 0.19 |
| MG63 | 1.30 ± 0.01 | KPD | 0.33 | MG63 | 4.00 | HOSMNNG | 0.32 |
| OS25HAL | 1.32 ± 0.01 | 143b | 0.59 | OS25HAL | 7.00 | U2OS | 0.39 |
| SJSA/OSA | 1.33 ± 0.04 | OS25HAL | 0.86 | HOSMNNG | 14.00 | OS25HAL | 0.54 |
| HOS | 1.83 | CAL72 | 1.33 | G292 | 16.00 | 143b | 0.60 |
| HOSMNNG | 3.03 ± 0.03 | MHM | 1.46 | SJSA/OSA | 25.00 | KPD | 0.75 |
| CAL72 | § | HOSMNNG | 1.48 | 143b | 43.00 | G292 | § |
| MHM | § | G292 | 1.72 | MHM | 55.00 | CAL72 | § |
| U2OS | § | U2OS | 1.74 | U2OS | 77.00 | MHM | § |
| 143B | § | SJSA/OSA | 1.79 | CAL72 | § | SJSA/OSA | § |
| Median | n.d | Median | 1.33 | Median | n.d. | Median | n.d |
| (BRCA-) |  | (BRCA-) |  | (BRCA-) |  | (BRCA-) |  |
| CAPAN1 | 1.41 ± 0.29 | CAPAN1 | 0.21 | CAPAN1 | 3.00 | CAPAN1 | 0.19 |

**Supplementary Table 2:** Inhibitory constant 50 (IC50) values in CAL72 RB1 normal OS with and with shRNA-mediated RB1 ablation. Data are based on inhibition of clonogenic activity. § denotes situations where an IC50 values could not be extrapolated from the concentration range assessed.

|  | Olaparib | Niraparib | Cisplatin |
| --- | --- | --- | --- |
| Cell line | IC50 (μM) | IC50 (μM) | IC50 (μM) |
| CAL72 unmod | § | 2.055 | § |
| CAL72 vector | § | 1.686 | § |
| CAL72 shRB1-1 | 0.408 ± 0.28 | 0.270 | 0.129 |
| CAL72 shRB1-2 | 0.563 ± 0.23 | 0.175 | 0.193 |
| CAL72 shRB1-3 | 0.493 | 0.432 | 0.076 |

## SUPPLEMENTARY MATERIAL AND METHODS

### Cell lines, chemicals and antibodies

STR profiling using the GenePrint 10 system (Promega) was used to confirm the identity of cell lines. All cell lines were maintained at 37°C in a humidified incubator with 5% CO2. Cisplatin and PARP inhibitors olaparib, niraparib, veliparib and talazoparib were purchased from Selleck Chemicals, dissolved in DMSO as a 10mM stock solution and stored at −80°C. Thymidine (Sigma), was dissolved in phosphate buffered saline (PBS) and used at a final concentration of 2 mM. For immunoblotting the following antibodies were used: anti-RB1 (4H1 clone), anti-phospho-CHK1 (Ser345), anti-phospho-CHK2 (Thr68) (C13C1), anti-CHK1 (2G1D5), anti-CHK2 (1C12) were purchased from Cell Signalling Technology, anti-GAPDH (6C5) was obtained from Abcam. For immunofluorescence analysis, anti-RAD51 (Abcam) and anti-phosphorylated histone-H2AX (Merck Millipore) antibodies were used.

### Clonogenic survival assays

Assays were performed as previously described (1). Cell were seeded in 6-well plates at a concentration of 1000-2000 cells per well and on the following day treated with vehicle or inhibitors. Two independent wells were used per conditions. After 10 to 14 days, plates were washed with Phosphate Buffered Saline (PBS), cell layers fixed with 4% formaldehyde for 10 minutes and stained for a minimum of 20 minutes using 0.025% f. c. crystal violet dye in PBS. Excess dye was rinsed off using repeat immersion in tap water. Plates were air dried and images captured using a desktop scanner. To quantify colony formation proficiency bound crystal violet dye was extracted using 30% v/v acetic acid, 30% v/v DMSO and 0.1% SDS and the optical absorbance determined spectrophotometrically at 570 nm using a multi-functional microplate reader. Clonogenic activity for inhibitor treated cells is expressed relative to that of vehicle-treated wells. Concentration-response curves were plotted using a three-parameter regression fit in GraphPad Prism 8.

### Immunofluorescence

Cells were seeded on glass coverslips placed on 6-well plates. 24 hours after plating cells were treated with inhibitors or ionising radiation. For analysis, cells were fixed in 4% paraformaldehyde in PBS for 10 minutes, washed twice with PBS, permeabilised with 0.2% Triton X-100 in PBS for 10 minutes, and incubated in 2% BSA in PBS for 20 minutes. Cells were then incubated at room temperature for 2 hours with primary antibodies diluted according to manufacturer’s recommendation in 2% BSA in PBS. Cells were washes in three changes of PBS followed by incubation with ALEXA Fluor 488 or 568 coupled secondary antibodies, diluted in 2% BSA in PBS for 1 hour. Cells were washed in three changes of PBS and mounted using VECTASHIELD™ (Vector laboratories) mounting medium containing DAPI. Cells were visualised using an LSM 510 Zeiss confocal laser-scanning microscope using the 63 × 1.4 NA oil objective. Z stacks were taken of cells representative and Maximum Intensity projections generated. For quantitative analysis, ≥ 100 cells from each condition were chosen at random and nuclear foci were counted manually to determine the percent positive for RAD51. γH2AX fluorescence was quantified using Imaris Cell Imaging software (Imaris), where at least 100 nuclei were analysed per experiment and condition.

### Immunoblotting

Samples were diluted to contain equal protein concentration prior to analysis. Protein concentrations in samples were measured by BCA protein assay (Pierce). Samples were separated on SDS polyacrylamide gels and transferred to an Immobilon-FL membrane (Millipore), before incubation with primary antibodies overnight at 4 °C. Membranes were washed with TBST (25 mM Tris, 140 mM NaCl, 0.1% Tween-20, pH 7.5) and probed with IRDye 680- or IRDye 800CW-conjugated (LI-COR), or HRP-conjugated secondary antibodies for 1 hour at room temperature.

The imaging and quantification of signals was carried out using an Odyssey CLx infrared imaging system or chemiluminescent detection (Pierce).

### Time resolved cell death analysis

Cells were seeded the day before the experiment at 10000-20000 cells per well of a 96 well plate or 40000 cells per well of a 12 well plate. The following day cells were treated with inhibitors at the indicated doses in the presence of 1□M SYTOX™ Green (Thermofisher) or 0.1□M Propidium Iodine (Sigma). Cells were imaged in real-time for the incorporation in 2 hours intervals using an IncuCyte ZOOM system, (essenbiosciences), as described before (1). Data were collected from a minimum of three parallel wells per condition and are typical for at least three independent assessments.

### Day-5 viability assessment using resazurin

Resazurin reduction assays were used to quantify the effect on inhibitor treatment cell survival and proliferation. Briefly, cells were seeded (1000-4000 cells/well) at densities optimised for each cell line in triplicates on clear, flat bottom 96-well plates (Corning). The following day, cells were treated with increasing concentrations of inhibitors at the indicated doses for 5 days added to wells as a 10-fold concentrated working stock, generated in cell culture media. Assessments were run in triplicate. At the end of treatment cells were incubated with resazurin (sigma) was added to a final concentration of 0.5 mM. Cells were incubated for 4 hours at 37°C to then read in a BioTek Synergy HT plate reader (excitation = 560 nm, emission = 590 nm). Mean Values derived from parallel triplicate wells were normalized to the mean value obtained for triplicate cell-free media control wells. Mean values for vehicle-treated control wells were assigned 100% viability and cell viability for inhibitor exposed wells calculated accordingly.

### Short hairpin constructs and viral infection

Viruses were packaged using HEK 293T cells. Cells were seeded into 10 cm tissue culture dishes for 24-36 hours, then transfected with packaging plasmids psPAX2, pMD2.G and lentiviral backbone construct at a ratio of 2.5:1:5.5 using the ProFection™ mammalian transfection system according to the manufacturer’s instructions (Promega). Transfection media was replaced with culture medium after 16 h incubation. Viral supernatants were harvested 24, 48 and 72 h later, filtered through 0.45-micron syringe filters, supplemented with hexadimethrine bromide to yield a final concentration 4 μg/ml (polybrene, Sigma-Aldrich) and either used directly or stored at −80°C. The pGIPZ-shRNA constructs targeting human RB1 were purchased from Horizon and were designated V2LHS 130606, V2LHS 130608 and V2LHS 340824 with sequences TAAGTTCACATGTCCTTTC, TTAACTGAAATGAAATCAC and AATCTTGCATCTAGATCTT respectively. For viral transduction cells were seeded into 6-well plates were incubated with polybrene supplemented viral supernatant in complete media overnight. The next day, media replaced with fresh media. Cells were subjected to puromycin selection 48 hrs post infection to select for virus uptake.

### Flow cytometry assisted cell cycle analysis

Cells were plated in 6-well dishes at a concentration of 1.5 x10^5^ cells per well and treated in parallel to cells seeded for experiments to document reposes to thymidine treatment. Thymidine (Sigma), dissolved in Phosphate Buffered Saline (PBS), was used at a final concentration of 2 mM. Cells were harvested eight hours later, fixed using 70% ethanol for a minimum of 12 hours. For analysis cells were treated with 10 □g/ml DNase and protease-free RNAse A for 10 minutes at room temperature, followed by addition of propidium iodide (Sigma) to a final concentration of 200 μg/ml containing. Samples were analysed using a Fortessa X20 (Becton Dickinson) flow cytometer.

### Human xenograft models

Breeder pairs of NRG (NOD.Cg-Rag1tm1Mom Il2rgtm1Wjl/Szj) immunodeficient mice were from the Charles River, UK. OHSN osteosarcoma-derived cells resuspended at a concentration of 8 × 10^7^ cells/ ml in sterile PBS and 50 μl (4X 10^6^ cells) were injected s.c. into the right flanks of host animals. Once tumours reached ~100 mm^3^, assessed using digital callipers, mice were randomly assigned to treatment groups and treated once daily p.o. talazoparib at 0.33 mg/ kg made up in using 10% Dimethylacetamide, 6% Solutol, and 84% PBS. Drug will be administered at 0.33 mg/ kg or vehicle, administered p. o. for three weeks, on 5 consecutive days (Monday to Friday) each. Mice were weighed, and tumour dimensions were determined twice weekly using digital callipers. Tumour volume was calculated using (length × width^2^)/2. Tumour growth was assessed relative to its respective initial size. All experiments were conducted using humane endpoints in accordance with AWERB guidelines.

### Mutational signature analysis

For mutational signature analysis in cell lines mutational data for nine of the cell lines derived from osteosarcoma were downloaded from the Broad Institute’s Cancer Cell Line Encyclopedia (**https://portals.broadinstitute.org/ccle**). Counts of the 96 single-base substitution triplet contexts were generated using sigProfilerMatrixGenerator (2). Each sample was decomposed into exposures of 46 previously published mutational signatures using sigProfilerSingleSample with default settings (2). Activities of all signatures identified in a sample were normalized in sum to 1. For mutational signature analysis in tumour sample, mutational data, mutational signature activities, and copy number data from a series of 37 previously published whole genome sequenced osteosarcomas was downloaded (3). Mutational signature activities were normalized in sum to 1 in each sample. Homozygous deletions, CN= {0,0}, and loss-of-heterozygosity events, CN= {>0,0}, were called at *RB1* from the associated copy number data. Combinations of mutations/copy number alterations of *RB1* were compared to COSMIC3 activity.

### Statistical Analysis

Two-tailed Student’s t test were used for normally distributed data and Mann-Whitney nonparametric test for skewed data that deviate from normality. Dunn’s multiple comparisons test was used for Immunofluorescence assay to correct for multiple testing errors. Survival was analysed by Kaplan-Meier plot, and log-rank (Mantel-Cox) test was used to compare data.

